# Genic Position and Methylation Context Shape DNA Methylation-Expression Relationships in Rice Internode Development

**DOI:** 10.64898/2026.07.09.737558

**Authors:** Niharika Nonavinakere Chandrakanth, Matthew T. McGowan, Nicolás Gaitán, Fan Lin, Vivian Ng, Anna Lipzen, Vasanth Singan, Chris Daum, Yuko Yoshinaga, Su Li, Lingtao Su, Dong Xu, Stephen P. Ficklin, Jorge Duitama, Laura E. Bartley

## Abstract

Elongating rice internodes present a developmental gradient from dividing meristem to mature cells, providing an elegant pseudo-time course for study of plant vegetative development. We tested the hypothesis that DNA methylation regulates gene expression during rice internode development by integrating RNA-seq and bisulfite DNA sequencing across eight internode segments. Previously described topologically associated chromatin domain borders aligned with transcription start sites of constitutive expressed genes. CpG and CHG differential methylation was enriched in young segments, consistent with maintenance methylation; whereas CHH methylation showed similar differential abundance in young and old segments. CHH and CHG methylation in upstream regions, CpG methylation within gene bodies, and any methylation in 5′ and 3′ untranslated regions were permissive of moderate to high gene expression. Very low expression was associated with CpG methylation upstream, CHG and CHH methylation within gene bodies, and CpG and CHG methylation downstream. A nonrandom subset of genes, including cell wall-related glycoside hydrolases, lignin and tricin biosynthesis enzymes, and WD40 proteins, showed methylation-expression correlations, with expression changes enriched in triple-marked elements. These results suggest that internode phenotypes of DNA methylation machinery mutants relate to alteration of specific target genes, opening approaches for grass culm improvement for lodging resistance and biomass production.

**Graphical Abstract:** 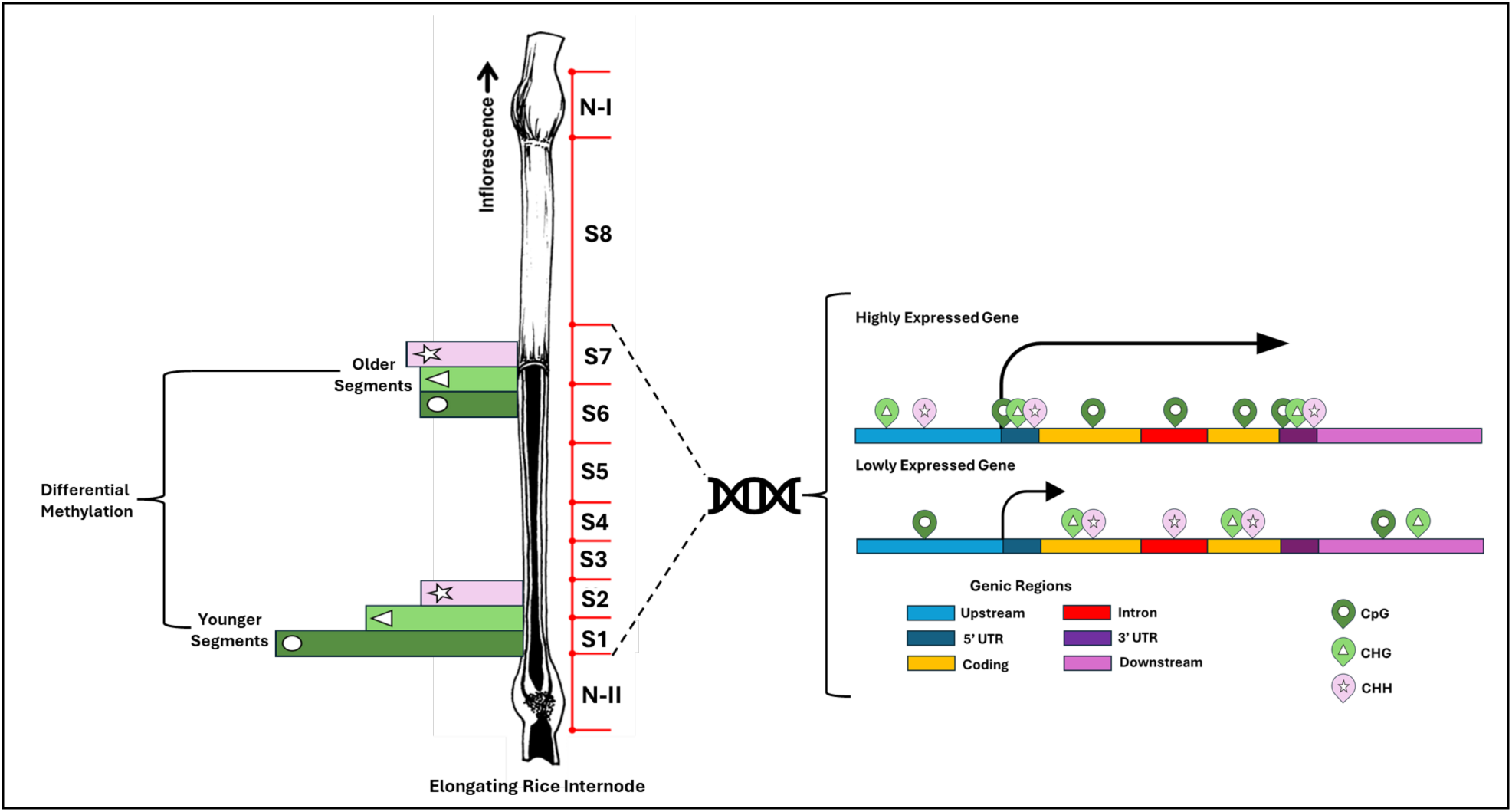

**Significance statement:** Stems of cereals and other grasses have an important function in supporting grain production and serving as a source of carbon for the bioeconomy. This study supports methylation of the cytosine bases of DNA as a regulator of chromatin structure and gene expression in the stems of rice. Different methylation contexts are implicated in specific roles in both positive and negative gene expression regulation that might be leveraged to improve stem properties.

## Introduction

In rice and other grasses, stems (i.e., culms) are composed of nodes and internodes that provide structural support and transport water and nutrients (Yano *et al*., 2015; Wang *et al*., 2018). A low stem-to-grain ratio was a driver of the green revolution (Gaur *et al*., 2020) improving lodging resistance (Kashiwagi *et al*., 2008), while more biomass is favored for biomass production for biorefining applications (Yang & Hwa, 2008; Le *et al*., 2022). Studied at the transcriptional level in multiple cereal species (Chen *et al*., 2013; Hirano *et al*., 2013; Patil *et al*., 2019; Xie *et al*., 2022), the grass internode during elongation represents a pseudo-time course in which different physical regions represent a progression of tissue development, simultaneously encapsulating the stages of cell division at the intercalary meristem, through elongation, to cellular maturation, culminating in secondary cell wall formation. Unlike other plant organs, such as roots and leaves, whose development also includes alterations in function (i.e., nutrient transport and photosynthesis), the internode primarily is a structural and transport organ throughout its development, potentially enabling investigation of developmental regulation without the confounding influence of shifting physiological roles.

In eukaryotes, DNA methylation is a crucial epigenetic signal, regulating genome stability, chromosomal structure, trait inheritance, and gene expression (Candaele *et al*., 2014; Han *et al*., 2018; Yin *et al*., 2024). Methylation primarily occurs at the fifth carbon of cytosine (i.e., 5-mC) in three sequence contexts: symmetric CpG and CHG, and asymmetric CHH (where H represents A, C, or T) (Kumar & Mohapatra, 2021).

Methylation in all three contexts (Zhong *et al*., 2021), but especially CpG and CHG methylation, is critical for genome stability, showing enrichment in heterochromatic regions characterized by transposable elements (TEs, repetitive DNA sequences that can move from one genomic location to another). TE-rich heterochromatin is associated with chromatin compaction and transcriptional repression (Zhong *et al*., 2021). Active and inactive chromatin segregates into A and B compartments at broad scales and self-interacting, topologically associated domains (TADs) at finer scales(Domb *et al*., 2022). Though appearing to have different functions in plants and animals (Santos *et al*., 2020), a commonality is that TAD interiors correspond to relatively compact chromatin environments, while TAD boundaries provide a more permissive context for transcription. Due to their symmetry, CpG and CHG methylation can be preserved by maintenance-related processes on hemi-methylated DNA after replication, providing long-term epigenetic memory that enables plants to inherit stable gene expression profiles (Jullien *et al*., 2006; Pikaard & Mittelsten Scheid, 2014). Both CpG and CHG are heritable, though they may be revised during reproductive transitions and to adapt gene expression to developmental needs (Bartels *et al*., 2018; Zhang *et al*., 2018; Tirot *et al*., 2021).

In contrast to the other marks, CHH methylation is both less abundant and qualitatively more dynamic, enabling environmental responsiveness and tissue-specific regulation (Kenchanmane Raju *et al*., 2019; Lee *et al*., 2023). In all three contexts, methylation of non-repetitive, non-TE regions protein-coding and non-protein-coding genes and regulatory sequences is also prevalent (Pikaard & Mittelsten Scheid, 2014; Bewick & Schmitz, 2017; Zhang *et al*., 2018). Methylation of non-TE genes can be repressive, blocking transcription factor binding or promoting chromatin inaccessibility, but methylation can also be stimulatory, as specific transcription factor complexes may bind more stably to methylated sequences (Héberlé & Bardet, 2019). *De novo* methylation processes add new methylation marks to previously unmethylated DNA sequences through the small RNA-directed DNA Methylation (RdDM) pathway (Markulin *et al*., 2021). This processes, which primarily targets CHH methylation, but can also alter CHG and CpG in specific contexts(Wang & Baulcombe, 2020), is essential for gene regulation under stress (Secco *et al*., 2015).

Alterations in DNA methylation accompany and alter stress responses and developmental changes. Drought stress induces both hypermethylation and hypomethylation at stress-responsive genes, modulating rice gene expression and stress adaptation (Garg *et al*., 2015; Wang *et al*., 2016; Wang *et al*., 2020). Genome-wide decrease in DNA methylation accompany fruit ripening (Ji & Wang, 2023). Mutations that alter DNA methylation (writing), DNA methylation recognition (reading), or DNA demethylation (erasing) influence rice stem growth and plant height, traits linked to internode elongation (**Tabel S1**). Individual disruption of CpG maintenance (MET1), CHG maintenance (CMT3), RdDM-mediated CHH deposition (AGO4/RDR2/DCL3), or active demethylation (ROS1) all result in reduced internode stature (Moritoh *et al*., 2012; Hu *et al*., 2021). In addition, perturbation of RdRM pathways alter hormone pathways involved in stem elongation (Wei *et al*., 2014). With respect to chromatin TADs, plants lack animal insulator proteins that define TAD boundaries (Acemel *et al*., 2017). Rather, rice TADs may be enriched in active chromatin features and, particularly, TCP (TEOSINTE BRANCHED1/CYCLOIDEA/PCF)-associated motifs (Liu *et al*., 2017). Functions of this class of plant-specific transcription factor include both positive (e.g., OsPCF7;(Li *et al*., 2020)) and negative (e.g., OsTCP4;(Ruan *et al*., 2026)) regulation of internode elongation. These observations suggest that methylation dynamics are required for normal stem growth and potentially represent an important and understudied aspect of vegetative plant development beyond a general effect of increased TE activity causing genome instability and hampering cellular function. Consistent with this, Candaele *et al*. (2014) showed that differential methylation in gene regulatory regions plays a role in the transition from cell division to cell expansion across four segments of the maize leaf developmental gradient, a system similar to the internode segments examined here.

Here, we present transcriptome and DNA 5-mC evidence for the actively elongating rice internode II supporting the hypothesis that DNA methylation is not a static epigenetic feature but rather a regulator of internode gene expression and development. CpG and CHG differential methylation were more abundant in early developmental stages, consistent with maintenance methylation in dividing cells, whereas CHH methylation showed similar differential abundance across young and mature segments. Methylation in different genomic contexts showed associations with gene expression, with both positive and negative correlations observed in TE and non-TE genes. The observation that specific gene categories displayed significant methylation-expression relationships provides evidence that the expression-methylation correlations are consequential for internode development.

## Materials and Methods

An extended methods is available in **Supplementary Methods 1**.

### Plant sampling

*Oryza sativa* ssp. *japonica* cv. Kitaake plants were grown under greenhouse conditions with natural light supplemented to maintain a 13-hour day length as described (Lin *et al*., 2017). To focus on development rather than environmental variation, internode sample collection at the booting stage occurred in multiple phases. Three replicates for RNA and then for DNA were harvested from groups of plants over the course of a few months between 13:00 and sunset. Internodes measuring 7.5–10.5 cm were dissected into up to 12 sections (**Figure 1A**). Node samples were excised at the swelling boundaries. Segments 1 to 4 were 5 mm in length, with segment 1 beginning immediately above the basal node. Segments 5 to 7 were 10 mm in length and Segment 8 consisting of the remainder before Node II. Some segment 1 samples were evenly bisected to further separate mature basal tissue (S1.1) from the youngest tissues (S1.2).

**Figure 1.**
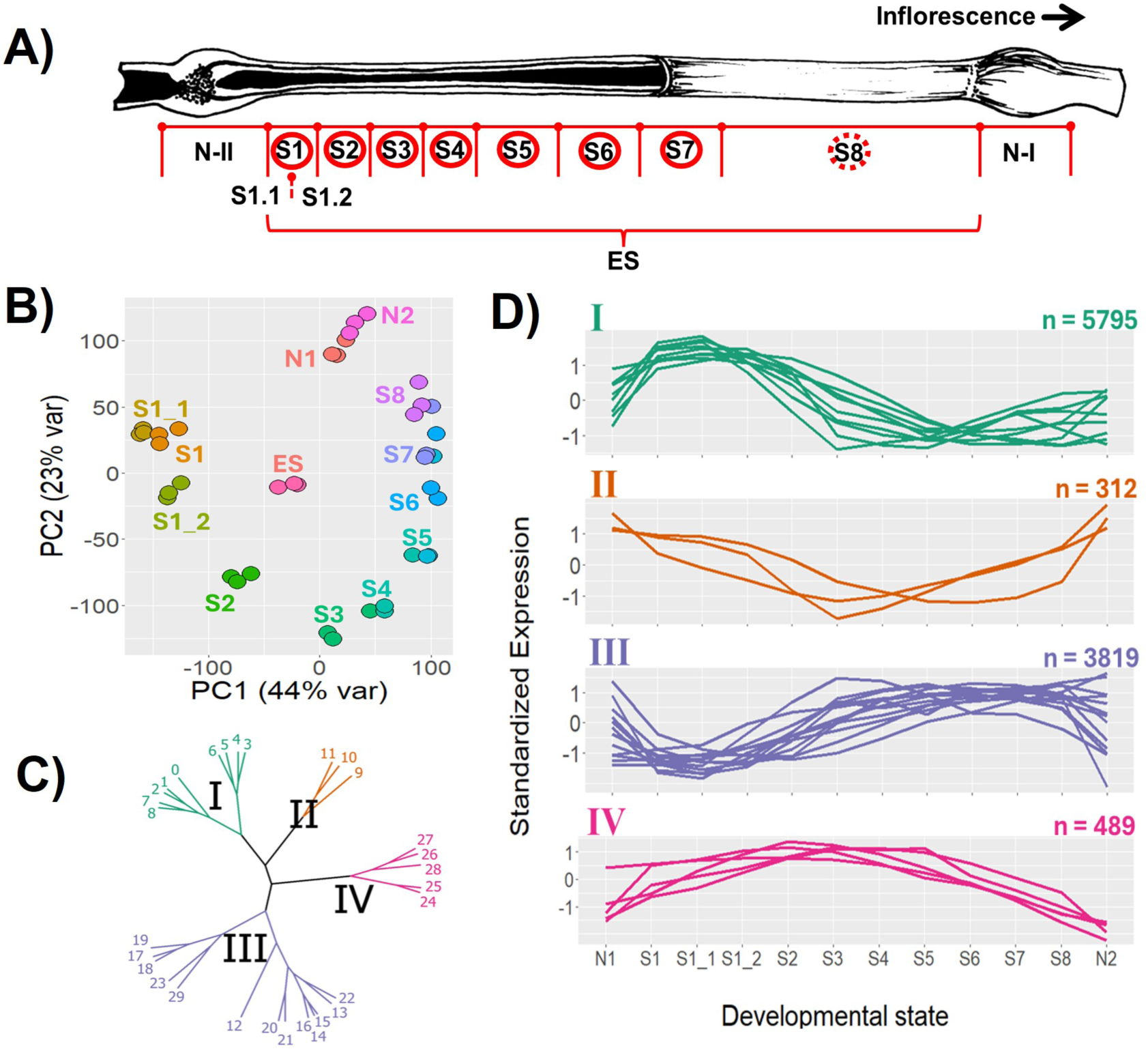
Developmental segmentation and transcriptome variation across the elongating rice internode. **(A)** The rice culm internode II and adjacent nodes were asymmetrically sectioned into twelve distinct segments for transcriptomics, labeled with black text. These segments consist of node (N-II), followed by the intercalary meristem-containing region (S1), which was bisected into subsections (S1.1, likely corresponding to the internode II “foot” (Tsuda *et al*., 2023), and S1.2), a transition zone (S2), elongation zones (S3–S4), and late maturation zones (S5–S8), terminating in the node (N-I) beneath internode I (the peduncle). Harvested segment lengths were as follows: S1 – S4, 5 mm; S5-S7, 10 mm; S8 2.5 to 5.5 cm. ES represents the elongating stem without nodes. DNA 5-methyl-C analysis was generated for segments S1-S8 (red circles); however, the S8 dataset (dashed circle) is excluded due to large variation. **(B)** Principal component (PC) analysis of transcriptome data from segments of the rice elongating internode. Each point represents a biological replicate, color-coded by segment. **(C)** Hierarchical clustering dendrogram of 30 co-expression clusters identified using CLUST (C0 - C29) shows distinct mega clusters (I–IV), indicated by branch colors: green (I), orange (II), purple (III), and pink (IV). **(D)** Standardized expression profiles of genes within each meta-cluster across development. Each line represents the mean standardized expression pattern of genes in a specific cluster. The number of genes in each meta-cluster is indicated (n).

### RNA preparation, sequencing, and analysis

RNA was prepared from approximately 85 mg of ground stem material and libraries prepared with with poly-A selection of mRNA, followed by pooling and sequencing on the Illumina HiSeq 2500 sequencer with a 2x150 indexed run recipe. Gene expression was quantified from the RNA sequencing data using transcriptome pseudo-alignment with gene-isoform predictions from the MSU/UGA v7 Nipponbare reference genome (Kawahara *et al*., 2013) using the GEMmaker nextflow pipeline v2.0 (Hadish *et al*., 2022). Read count and TPM values were calculated at the isoform level using Kallisto (Bray *et al*., 2016) in bootstrap mode (-b = 100). Principal component analysis (PCA) was performed using base R functions with results visualized via ggplot2.

### Chromatin analysis

TAD border data from *Oryza sativa* previously identified using the Armatus algorithm (Liu *et al*., 2017; Golicz *et al*., 2020) were obtained and gene transcription start sites (TSS) used to calculate the distance to the nearest TAD border. To classify genes based on expression patterns across samples, the proportion of zero-expression values for each gene was calculated using the mean-normalized expression matrix. For each gene, the number of samples with an expression value of zero was divided by the total number of samples to obtain the zero-expression proportion (zeroprop). Genes were then categorized into three expression types: constitutive (zeroprop = 0); repressed, (zeroprop = 1), and semirepressed, (0 < zeroprop < 1). A Kolmogorov–Smirnov (KS) test indicated the non-normality of TAD distance distributions, and a Mann-Whitney U test compared TAD distances between non-TE and TE genes. To evaluate differences across the six groups defined by expression type and TE status, we employed a Kruskal-Wallis test, followed by Tukey’s post hoc test. P adjusted < 0.05 were considered significantly different.

### DNA preparation, bisulfite-sequencing collection, processing, and analysis

DNA was prepared using a Qiagen DNeasy Plant Mini column from ∼75 mg of frozen ground stem biomass. DNA was sheared to 600 bp, size selected, ligated with adaptors, and bisulfite treated, converting the non-methylated cytosine to uracil, followed by 10 cycles of PCR amplification. Libraries were multiplexed, pooled, and sequenced on an Illumina NovaSeq 6000 following a 2x150 indexed run recipe. Bismark v0.22.3 (Krueger & Andrews, 2011) generated a modified version of the Nipponbare MSU7 reference genome (Kawahara *et al*., 2013) and mapped reads for each sample. Raw counts of methylated and unmethylated base pairs were obtained for each alignment file running the methylation extractor. A custom script based on the Next Generation Sequencing Experience Platform (Tello *et al*., 2023) calculated the percentage of methylation across loci from each sample. Genic regions were as follows: coding exons, introns, 5’-UTRs, 3’-UTRs, upstream (300 bp before the 5’-UTR) and downstream (300 bp after the 3’-UTR). The total number of possible methylation sites was calculated for each region and context. For each sample, a site was considered if the raw depth (methylated and unmethylated calls) was at least 5. Each site passing the depth filter was called methylated if at least 30% of the calls support the methylated allele.

An R script was built to analyze the matrix of percentage of methylated base pairs generated for each context. Five samples (BS1S6, BS2S8, BS3S4, BS3S7 and BS3S8 (BS “Replicate Number”; S “Segment Number”)) with more than 25% of missing data were removed from the analysis. Regions having counts for less than 15 of the 19 retained samples were also removed. Missing data points still present in the dataset were imputed with the average of the non-missing data within the region. Finally, regions having a standard deviation of at least 5% were selected for sample clustering and for correlation with expression data.

Differential methylation was calculated as the average methylation percentage in the older segments (S6 and S7) minus the younger segments (S1 and S2) with ≥ 10% change in methylation and Fisher’s exact test P < 0.05. Thus, positive values indicate higher methylation in older segments, whereas negative values indicate higher methylation in younger segments. For each gene, this analysis considered the genic region with the highest methylation. Functional enrichment analysis used the comprehensive annotation of rice multi-omics data (CARMO) platform (Wang *et al*., 2015)..

### Transcriptome-DNA methylation analysis

For evaluation of genic regions associated with transcript abundance, transcriptome data were divided into very low and lowly expressed with a cut-off of VST < 7.5. Methylation effect sizes on gene expression were estimated as log₂ odds ratios from 2×2 contingency tables with a 0.5 continuity correction, and significance was assessed using Fisher’s exact test. P-values were adjusted for multiple testing using the Benjamini–Hochberg false discovery rate (FDR) procedure, and associations with P_adj_ < 0.05 were considered significant. For correlation analysis, the average over replicates of segments S1 to S7 was calculated. Genes with regions having Spearman correlation P < 0.05 were selected for functional enrichment analysis and clustering analysis using CARMO. For all statistical analyses, we employed the Kruskal-Wallis test, followed by Dunn’s post-hoc non-parametric test, due to the non-normal distribution of the data. Statistical significance was determined at a P < 0.05.

## Results

### Transcription patterns of internode development

To provide a context for methylation analysis, we performed RNA-Seq on twelve asymmetrically divided segments (S) of the elongating rice internode II of *Oryza sativa* ssp. *japonica* cv. Kitaake (**Figure 1A**). Of 39,045 non-TE annotated loci (Kawahara *et al*., 2013), 27,860 (71%) were expressed in the internode. Of 6,932 annotated TE loci, 4,621 (27%) gave detectable expression (**Figure S1**). Principal components analysis (PCA) revealed distinct, but continuous differences among segments, with good replicate agreement (**Figure 1B**).

Clustering of co-expression profiles using spatial ordering (N-I, S1, S1.1, S1.2, S2, S3, S4, S5, S6, S7, S8, N-II) identified 30 co-expression clusters (**Figure 1C, Figure S2**), encompassing 9,908 non-TE and 1,801 TE gene models. Hierarchical clustering identified four distinct meta-clusters: early-stage (I), early & late-stage (II), late-stage (III), and mid-stage (IV) expression (**Figure 1D**). The early-stage (I) and late-stage (III) meta-clusters contained the greatest numbers of genes, suggesting that transcriptional regulation is most dynamic between these developmental stages, with informative gene annotation enrichments (**Data S1**).

### Chromatin structure and gene expression

Due to the relationships between chromatin domains, gene expression, and DNA methylation (Zhang *et al*., 2019), we next investigated whether chromatin features of the rice genome could help explain transcriptional activity in the internode. Prior work indicates that structures such as TADs are somewhat conserved across tissues and genotypes (Golicz *et al*., 2020; Sun *et al*., 2020; McArthur & Capra, 2021). Using TADs identified in rice leaf (for *O. sativa* ssp. *japonica* cv. Nipponbare) samples (Liu *et al*., 2017), we stratified TE and non-TE genes into constitutive, semirepressed, and repressed groups based on internode expression. Constitutively expressed non-TE gene transcription start sites were centered at TAD borders, while semirepressed and repressed genes, regardless of TE status, were broadly distributed and shifted toward intra-TAD regions (**Figure 2**). A Kruskal-Wallis test indicated significant differences among the six expression-gene class combinations (P = 1E-269), and Dunn’s post hoc test with Holm correction showed that pairwise comparisons were significantly different except between semirepressed non-TE and TE genes and between repressed non-TE and TE genes (**Figure 2**). Thus, TAD stability within the *japonica* subspecies and across tissue types appears to be a factor in gene expression control.

**Figure 2:**
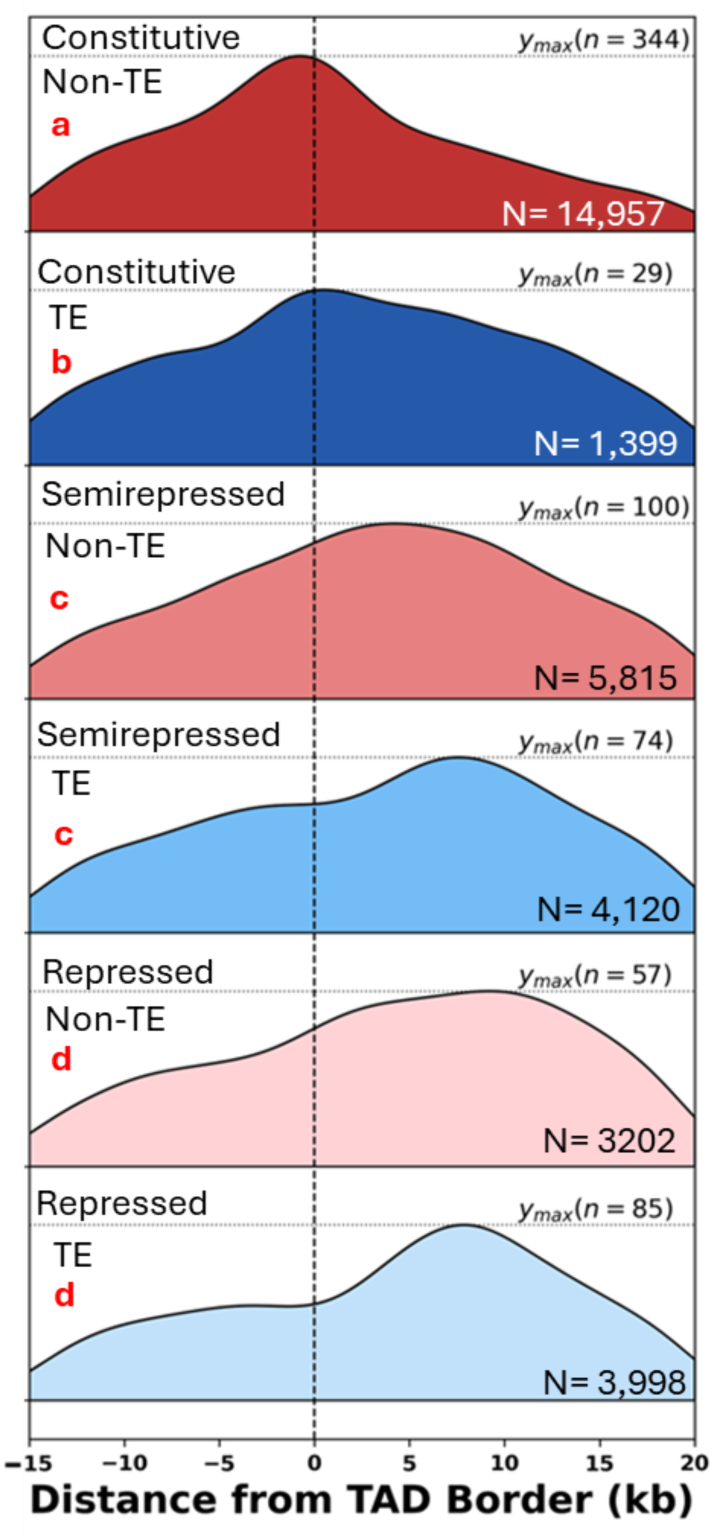
Constitutively expressed non-TE and TE genes are enriched near topologically associated domain (TAD) borders compared with semirepressed and repressed genes. Plots show the normalized density of genes as a function of distance from the nearest TAD border in kilobases (kb). Negative values indicate that transcription start sites (TSSs) are located outside TADs, whereas positive values indicate TSSs within TAD interiors. Genes are grouped by expression status (constitutive, semirepressed, and repressed) and gene class (non-TE or TE-associated). Non-TE genes are shown in reds, and TE-associated genes are shown in blues. The total number of genes in each category is indicated in each panel as N; y_max_ (n) values indicates the highest bin frequency. Red letters indicate Dunn’s post hoc groupings, where groups sharing the same letter are not significantly different at α = 0.05.

### DNA methylation patterns across internode development

To examine relationships between DNA methylation and internode gene expression, we used bisulfite sequencing to map cytosine methylation patterns in internode segments S1-S8 (**Figure 1A**). Bisulfite sequencing relies on conversion of unmethylated cytosines to uracil (Wreczycka *et al*., 2017), revealing the methylation status of cytosines in all CpG, CHG, and CHH contexts. Due to high variation, all S8 samples were removed from further analysis (**Figure S3**). To ensure high-confidence methylation detection, a minimum depth of 5 reads was required at each site, while a threshold of ≥30% methylated was applied to distinguish confident signals from background (See Methods). After filtering, PCA of methylation occurrence revealed low correlation of biological replicates, especially in early development (**Figure 3A**). PC2 explains ∼8% of the variance, and separates the early stages, S1-S3, from the latter ones, S4–S7. If of biological origin, this pattern suggests highly variable early-stage methylation that transitions to more similar profiles in later stages. After filtering, we detected approximately 128k methylation events in non-TEs and 26k in TEs. Eighty-three percent of loci showed robust evidence of methylation, and 45% were methylated in all three contexts (**Figure S1**). Aligning with previously reported rice results (Wang *et al*., 2020; Ni *et al*., 2021), CpG methylation was the most prevalent context in both non-TE and TE genes, followed by CHG and then CHH (**Tabel S2)**.

**Figure 3:**
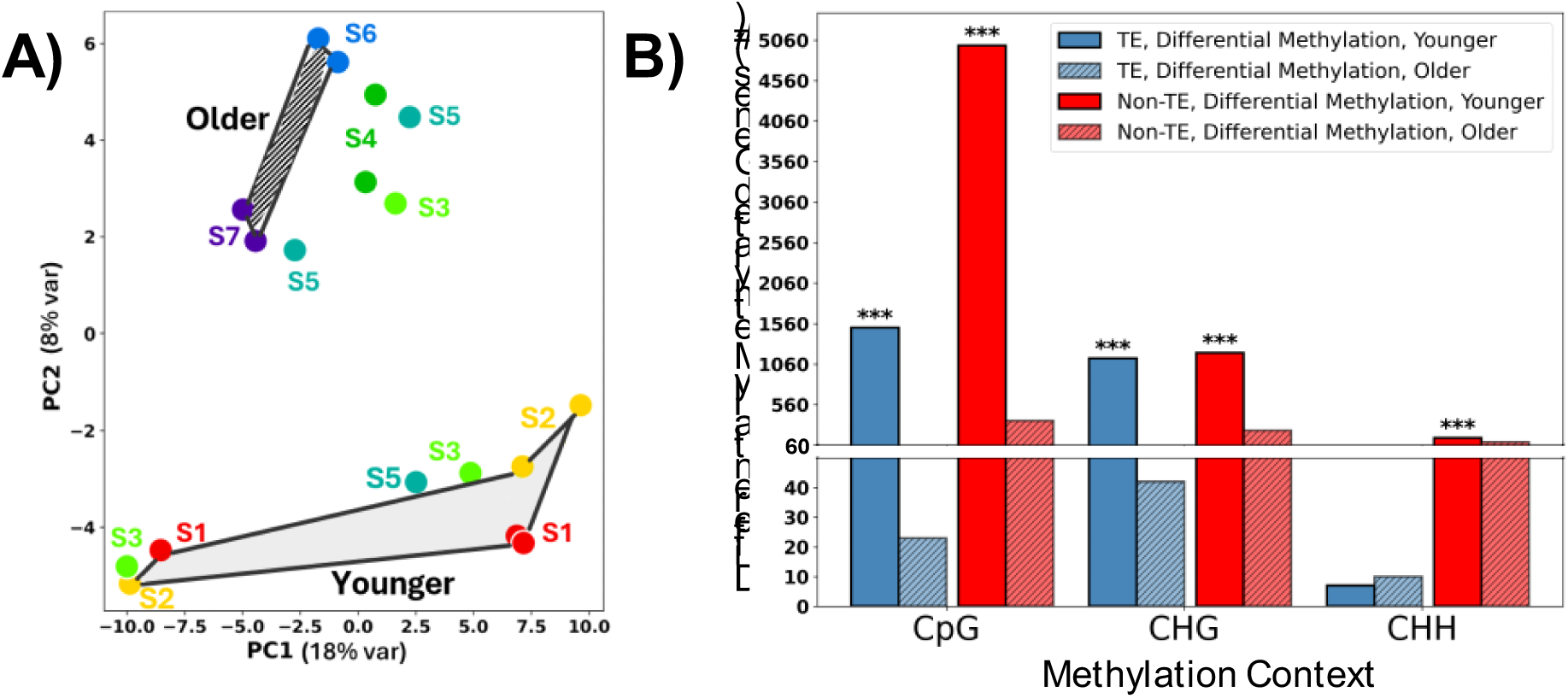
Dynamics of DNA methylation between early and late internode development. **(A)** Principal component (PC) analysis of methylation data across internode developmental stages. The plot shows PC1 and PC2 derived from methylation data across stages S1–S7. Each point represents a biological replicate, colored by stage. The shading and hatching denote the samples used in the comparison in B, representing young and old segments, respectively. **(B)** Differential methylation patterns in methylation contexts and segments. The chart illustrates the number of genes that exhibit significant differential methylation in younger (S1-S2) vs. older (S6-S7) segments across CpG, CHG, and CHH methylation contexts for TE (blue bars) and non-TE-associated (red bars) genes. Solid bars represent genes with higher methylation in younger segments. Hatched bars represent genes with higher methylation in older segments. In all contexts, younger segments (solid bars) show a significantly higher frequency of differential methylation. (***, Binomial test, P value < 0.001), but the fold effect is greatest in CpG and CHG contexts. Only genes with ≥10 percent change and statistical significance (Fisher’s exact test, P value < 0.05) are included in this analysis.

As expected, methylation frequency generally distinguished TE and non-TE genes. CpG and CHG methylation frequency (% methylated sites out of possible sites) was higher in TEs vs non-TE genes (75% CpG-TE vs. 60% CpG-non-TE, 65% CHG-TE vs. 30% CHG-non-TE; **Figure S4**). In contrast, CHH methylation was less frequent in TEs (4%) than in non-TE genes (18%), consistent with previous observations that CHH methylation plays a relatively minor role in TE repression and instead may modulate non-TE gene regulation. Methylation profiles of TEs and non-TEs remained largely consistent across internode segments, with only minor, though statistically significant global declines in CpG and CHG methylation between younger and older segments (Kruskal-Wallis-Dunn’s post-hoc, P < 0.05).

A subset of genes showed differential methylation (DM) between young and old segments, with distinct trends depending on methylation context. Differential methylation was calculated as the average methylation frequency difference between older (S6 and S7) and younger segments (S1 and S2). Genes with ≥ 10 percent differential methylation (DM, Fisher’s exact test, P < 0.05) were then categorized by whether methylation increased or decreased from young to old. For 8,459 out of 26,943 consistently methylated genes (31% of the 32,490 expressed loci - **Figure S1**), methylation varies significantly between early and late segments. Significantly higher CpG and CHG DM was observed in younger segments compared to older segments for both non-TEs and TEs (**Figure 3B**; Binomial test, P < 0.001). CHH methylation exhibited similar DM between young and old in both TE and non-TE genes, though the CHH non-TE DM genes were also statistically significantly more abundant in young segments. Serine/threonine protein kinases were strongly enriched among CpG and CHG DM genes in both younger and older segments (P_adj_ < 0.05). For CHH methylation, kinases were enriched only in older segments (P_adj_ < 0.05; **Data S2**).

### Integration of Methylation and Transcript Abundance in Internode Development

The position of methylation events within a gene can differentially affect gene expression (Zhang *et al*., 2006; Bewick & Schmitz, 2017). We calculated methylation percentages for each genic element (i.e., up- and down-stream, untranslated regions, coding sequence, introns). TE and non-TE genes showed differences in the uniformity of methylation across genic regions (**Figure S5**), but similar transcript abundance relationships with methylation context and genic position (**Figure 4, Table S3**). For this analysis, we divided genes into very low and permissive (i.e., moderate to high) expression categories based on the distributions of variance-stabilized transformed (VST) gene expression values (**Figure S4C and S4D**). Very lowly expressed genes (VST ≤ 7.5) were enriched for CpG in their up or down-stream regions and/or for CHG and CHH coding-region methylation (**Figure 4A & 4B**). In contrast, permissive expression (VST > 7.5) was associated with CHG and CHH upstream methylation, CpG coding sequence methylation, and both 5′-UTR and 3′-UTR methylation in any context (**Figure 4A & 4C**).

**Figure 4:**
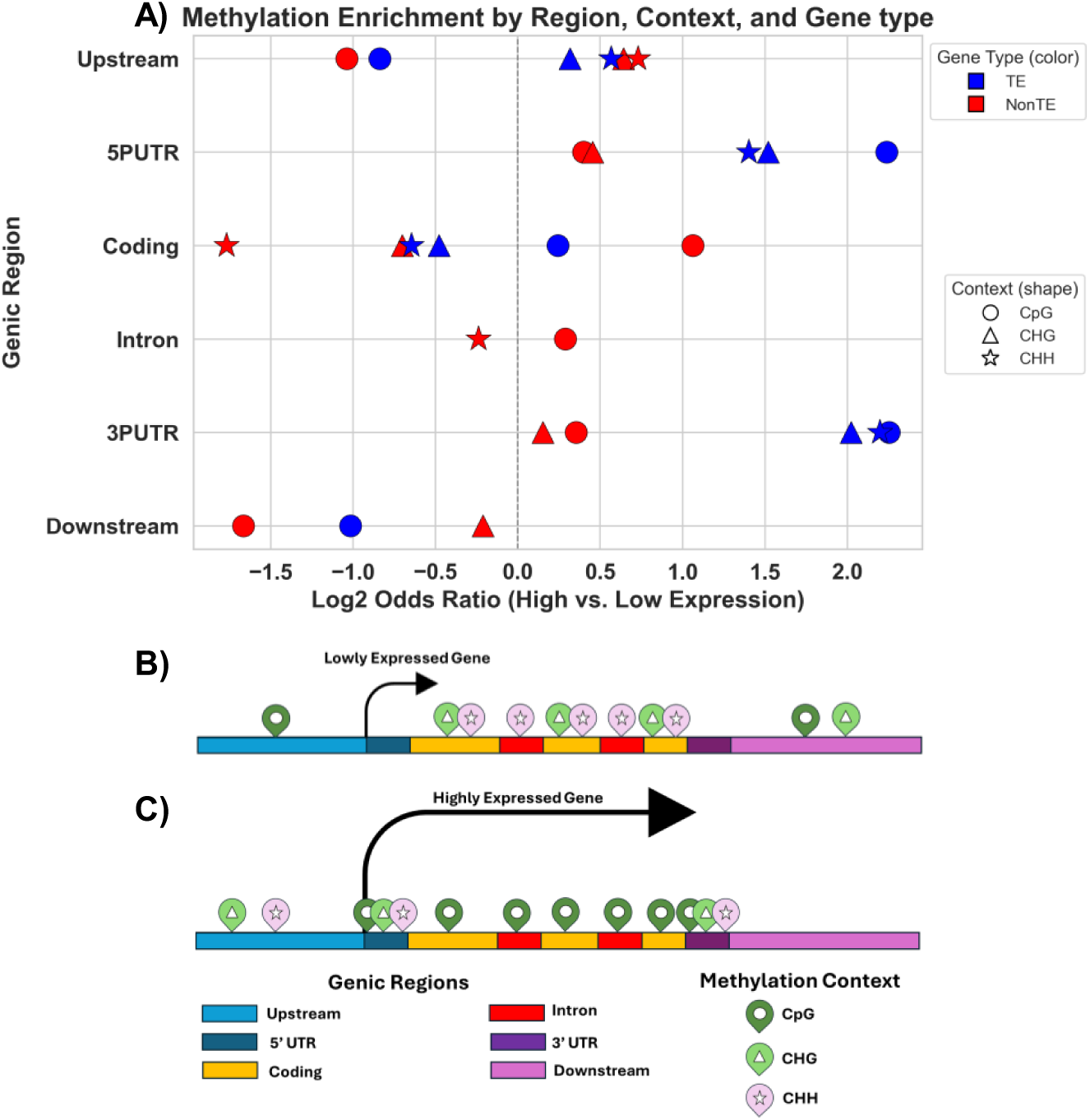
Significant associations between DNA methylation context and gene expression depending on genic regions. **(A)** Enrichment of DNA methylation for different genic regions in genes with moderate to high expression (variance stabilized transcript abundance, VST > 7.5) versus very low expression (VST ≤ 7.5). Each point represents a specific methylation context and gene class, with shapes indicating methylation context (CpG – circle, CHG – triangle, CHH – star) and color distinguishing gene class (TE – blue, non-TE - red). Positive values indicate enrichment in permissively expressed genes, and negative values indicate enrichment in very lowly expressed genes. Statistical significance for each comparison was assessed using Fisher’s exact test followed by Benjamini–Hochberg FDR correction, with all displayed points representing significant results (P_adj_ < 0.05). Summary by genic region of specific DNA methylation patterns associated with very low gene expression **(B)** and permissive gene expression **(C)**. Genic region is indicated by colored bars. CHG is indicated by light green and triangles, CHH by pink and stars, CpG by dark green and circles. Data for this figure are in **Table S3**.

### Correlations between methylation and non-TE transcription

Taking advantage of the gradient of samples in the internode pseudo-time course, correlation analysis between non-TE transcript abundance and methylation frequency revealed relationships enriched for particular methylation contexts (**Figure S6; Data S3**). Spearman correlation was prioritized as DNA methylation and transcription is often non-linear (Lou *et al*., 2014). **Figure 5A** shows the distribution of Spearman correlation coefficients for CpG, CHG, and CHH methylation classes, revealing that ∼12% of methylation marks were significantly correlated with transcript abundance. Both CpG and CHG included more significant negatively correlated than positively correlated genes. In contrast, CHH methylation was even between positive and negatively correlated genes (**Table S4**, **Data S3**). After Benjamini-Hochberg correction, no significant differences in genic positional distribution were detected between positively and negatively correlated genes (although minor unadjusted trends were observed in some regions, **Table S5; Figure S7**), so these were combined for further analysis (**Table S6**).

**Figure 5:**
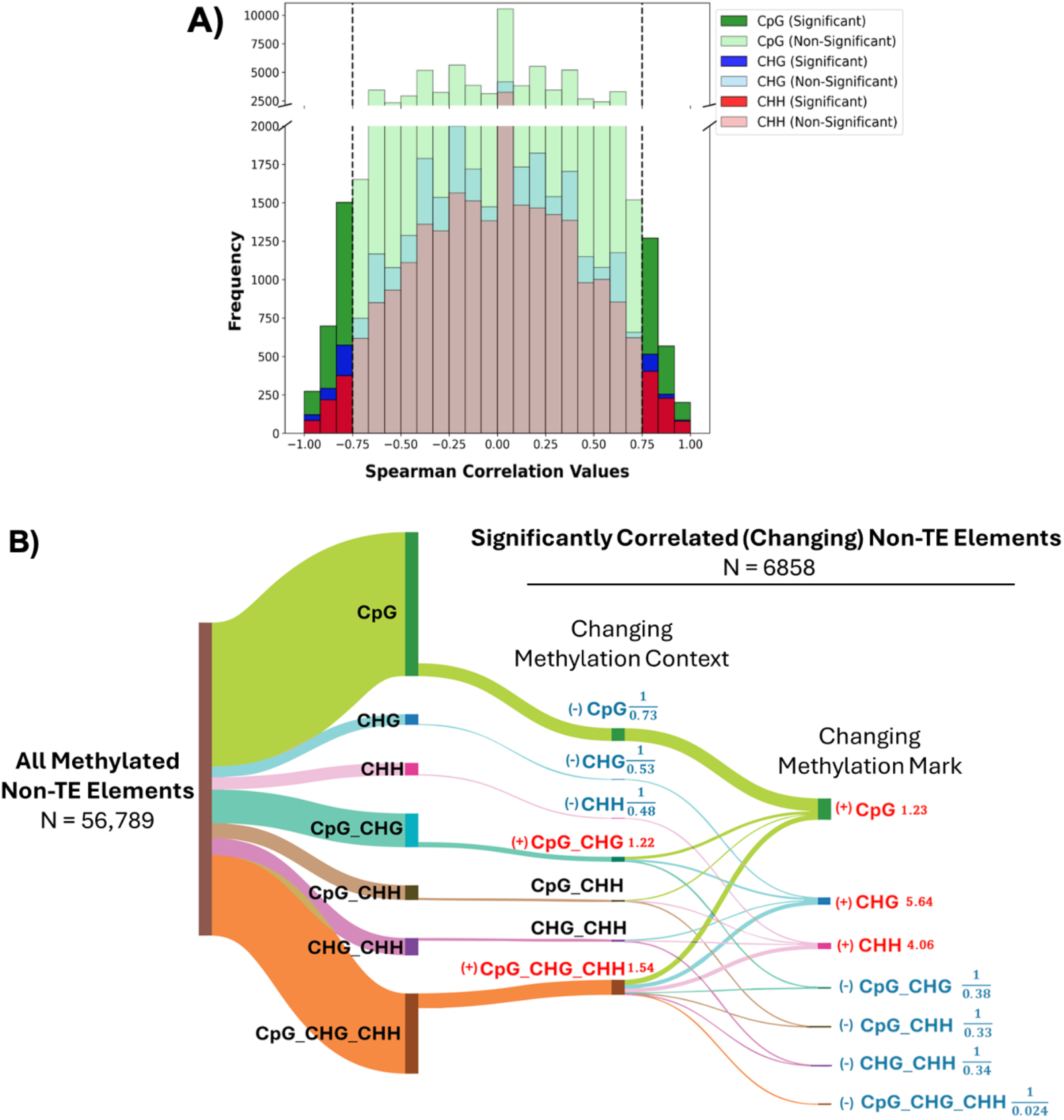
Significant DNA methylation-expression correlations of non-TE genes are more likely for single marks from genes with multiple marks. **(A)** Distribution of Spearman correlation values for CpG (green), CHG (blue), and CHH (red). Lighter bars are non-significant correlations, and darker shades are statistically significant (P < 0.05). **(B)** Distribution and transitions of DNA methylation across non-TE genes. The left side shows the global abundance of methylation contexts and their combinations across all methylated non-TE genic elements (N = 56,789). The middle section depicts the methylation context for the subset of elements for which methylation is significantly correlated with expression of the corresponding gene (N = 6858). The right panel displays the specific methylation marks that are changing within those contexts. Flow widths are proportion to number of elements. Fold-enrichment values (colored text) compare observed versus expected frequencies relative to the global methylation distribution; red text indicate significant enrichment, and blue text indicates significant depletion (hypergeometric test, P < 0.001).

**Figure 5B** traces associations of methylated contexts of non-TEs with the 12% of expression-correlated loci, highlighting that expression associations are typically driven by selective remodeling of single CpG, CHG, or CHH marks. Globally, loci exhibited a mixture of single and combined methylation contexts, with CpG-only, CpG_CHG double, and CpG_CHG_CHH triple marks being the most frequent categories (**Figure 5B; left panel**). A change in only single marks was likely to correlate with transcript abundance, with CpG over-represented by 1.2-fold, CHG by 5.6-fold, and CHH by 4.1-fold (hypergeometric test, P < 0.001; **Figure 5B, right panel**). Tracing the context of the methylation and expression-correlated loci revealed that the dynamic marks were disproportionately from CpG•CHG (1.2-fold enriched) and CpG•CHG•CHH (1.5-fold enriched) loci (**Figure 5B; middle panel**). On the other hand, correlations between methylation in any single methylation context and transcript abundance was observed about half as often as expected (**Figure 5B; middle panel**). Examining genic regions within the dataset failed to yield a robust signal due to the limited sample size along with multiple-testing correction (**Table S7**). Similarly, separate analysis of TE genes and their genic regions, revealed similar but not statistically significant trends (**Table S8)**. Together, these patterns support a model in which expression changes are preferentially associated with context-specific tuning of individual marks, especially CHG and CHH, at the subset of loci with a preference for methylation in multiple contexts, especially triple marks.

Functional category analysis of transcripts with significant methylation correlations revealed notable enrichments in specific biological processes, suggesting that methylation-transcript correlation relationships are non-random. **Table 1** summarizes statistically significant CARMO-enriched functional clusters by methylation context. Transcripts for glycoside hydrolases, which conduct cell wall modification and remodeling, were enriched among positively correlated genes across all three methylation contexts (P < 0.05, fold enrichment > 2, **Data S4**). WD40 domain-containing protein transcripts showed enrichment in both positive and negative CpG methylation correlations (**Table 1**), among which multiple loci have *A. thaliana* orthologs implicated in cell wall-related processes (Guerriero *et al*., 2015; Jain & Pandey, 2018). We also note that serine/threonine kinases were positively correlated with CHH methylation (P < 0.05; **Data S4**). Of particular relevance to the developing internode, literature curated lignin and tricin biosynthesis genes were significantly enriched for CHH-negative correlations within the set of methylated genes (P_adj_ = 2.E-04, **Data S5**).

**Table 1:**
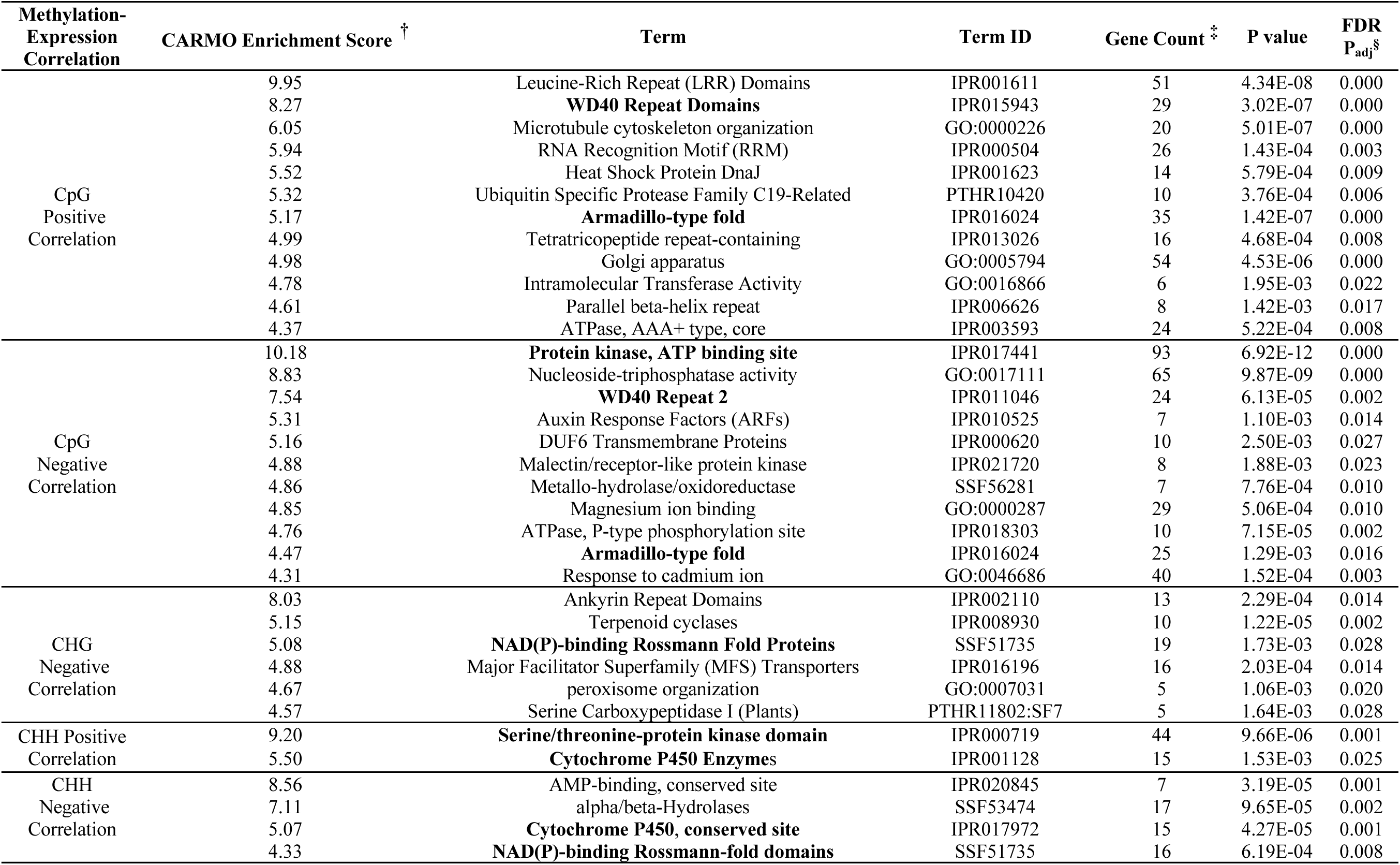
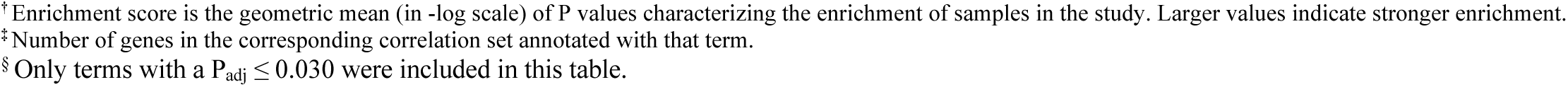
Top Comprehensive Annotation of Rice Multi-Omics data (CARMO) platform (Wang *et al*., 2015)-enriched gene clusters among significant positive and negative DNA methylation–expression correlated genes, P_adj_ < 0.05. Repeated terms are shown in bold. **Data S4** includes the complete enrichment results.

### Internode Expression Patterns of Methylation Machinery

To understand the proteins that might be driving the observed methylation patterns, we examined the transcriptional profiles of cytosine methylation writers, readers, erasers, and TCP proteins (**Table S9; Figure S8; Figure S9; Figure S10**). Genetically characterized 5-mC writers such as DNA Methyltransferase (MET1-2), which maintains CpG methylation, and Chromomethylase 3 (CMT3), responsible for CHG methylation, are part of meta-cluster I with higher expression in early segments. In contrast, CHH methylation regulators, such as Domains Rearranged Methylase 2 (DRM2) and Chromomethylase 2 (CMT2), are in meta-cluster III with higher expression in later segments (**Figure S8**). These results align with the greater abundance of CpG and CHG differentially methylated genes in young segments compared to the older segments and contrasting results with CHH (**Figure 3B**). Reader proteins, such as Methyl-CpG-Binding Domain (MBD) (Qu *et al*., 2021), SUVH (SU(VAR) 1-11 Homologs) (Qin *et al*., 2010), and SHH1 (SAWADEE HOMEODOMAIN HOMOLOG 1) (Law *et al*., 2011), exhibited diverse clustering profiles potentially related to their roles in recognizing methylation marks. While many of these proteins remain understudied, their expression patterns suggest distinct functional roles in different developmental stages. Eraser proteins, including Repressor of Silencing (e.g., ROS1a) and Demeter-Like (e.g., DML3) proteins (Lanciano & Mirouze, 2017), were predominantly expressed in mid-to-late internode segments (meta-clusters II, III, and IV), consistent with increased demethylation activity in later stages of internode development. Finally, of the 22 TCP genes described in rice (Yao *et al*., 2007), 18 are expressed in the internode transcriptome dataset, with distinct patterns across the segments (**Figure S9**).

## Discussion

Integration of transcriptome and global DNA methylation evidences regulatory associations between transcript abundance and methylation within the pseudo time course of internode development that potentially explain methylation mutant phenotypes (**Figure S11**) and suggest methylation roles that are difficult to study with mutants. The insights provide a foundation for understanding how DNA methylation coordinates developmental transitions and tissue-specific gene regulation in plants.

### Evidence for involvement of TAD domains in transcriptional regulation

Our findings support transcriptional control related to TAD chromatin organization being generally conserved in rice. Unlike in mammals (Valton & Dekker, 2016; Akdemir *et al*., 2020), TADs in plants lack a clear boundary marker, which is exacerbated by plant TAD identification being based on pattern-recognition algorithms the results of which depend on parameter values leading to ambiguity about the function of plant TADs (Santos *et al*., 2020). The consistency of relationships with gene expression that we observe provides further support for the notion that TADs are generally stable structures. Without direct measurement of TADs, we are unable to address subtle differences in structure, such as observed when comparing ear vs. tassel TADs (Sun *et al*., 2020). As plant genomes typically have high TE content, plant TADs may have evolved to play a particularly important role in TE repression. Our observation that TAD structure appears strongly associated with TE repression may be biased toward the most stable TAD configurations, which are likely to enforce TE silencing under diverse conditions. Supporting a role for chromatin structure in internode development, a recent study disrupting two nucleosome remodelers in rice reported severe internode developmental defects (Ikram *et al*., 2022). These results point to an interplay between TAD organization, nucleosome dynamics, and transcriptional regulation, with core TAD features conserved across cultivars, organs, and developmental stages in rice.

### Genic position and context-dependent methylation correlate with transcription

DNA methylation along with expression stratification and transcript abundance correlations are consistent with an active, specific role for methylation in internode gene expression. Stratification of transcript abundance revealed global, stable methylation patterns relate to permissive expression in internodes, with similar effects but different frequencies for TE and non-TE genes (**Figure 4**). CpG methylation, which is more frequent in TEs than non-TE genes (**Figure S5**), is repressive when occurring in both upstream and downstream genic regions, consistent with upstream regions of transposable elements being frequently methylated and transcriptionally repressed (Berdasco *et al*., 2008; Lister *et al*., 2008), as observed in rice internodes (Wang *et al*., 2020). Likewise, CpG is depleted in Arabidopsis accessible chromatin regions both in TE and non-TE genes (Sullivan *et al*., 2014).

On the other hand, other methylation marks, particularly upstream CHH and CHG, gene body CpG methylation, and any methylation in UTRs is permissive of gene expression for both TEs and non-TEs. Arabidopsis accessible chromatin regions have also been observed to be enriched for CHH methylation (Sullivan *et al*., 2014). Similarly, analysis across eight grass genomes found that mCHH “islands” enriched near genes, often within ∼2 kb upstream of the TSS and show a generally positive but weak association with gene expression (Martin *et al*., 2021). Transcriptional permissiveness due to CHG upstream methylation has been noted less in the literature, though was observed previously in rice internodes (Wang *et al*., 2020). A positive association between CpG gene body methylation and transcription has also been observed (Wang *et al*., 2020), and some evidence suggests that it is under selective pressure, though the mechanism of gene expression control remains an area of active study (Muyle *et al*., 2022).

Correlation analysis between non-TE transcript abundance and methylation provides evidence that methylation context matters for methylation control of development (**Figure 5B**). Increased or decreased methylation, especially CHG and CHH in the context of double (CpG•CHG) or triple methylation marks, correlate either positively or negatively with gene expression. Thus, addition or removal of methylation within specific methylation contexts is either more likely during or causes transcriptional changes. The importance of context was illustrated by recent work suggesting that sequence impacts CpG demethylation and transcriptional stimulation (Stefansson *et al*., 2024); likewise, rice CMT3b functions in methylation in CG-rich regions (Hu *et al*., 2021). Though work is needed to better understand the importance of these observations, specific gene families are enriched for methylation-transcript correlations supports a model of methylation alterations as a mechanism of large-scale transcriptional regulation during plant development.

### Enrichments in differentially methylated and transcript abundance-DNA methylation correlated genes

CARMO-enriched functional clusters of DM genes and transcripts that correlate with methylation highlight potential roles for methylation for control of particular gene classes (**Table 1**). Each methylation class and correlation sign shows enrichment in specific functional categories, and several enriched annotations are shared among methylation contexts. For positively correlated genes, methylation may stimulate expression or demethylation may repress expression, while for negatively correlated genes, methylation may repress expression or demethylation may be stimulatory.

Repeatedly enriched terms among methylation-transcript abundance correlated genes include signaling and protein-interaction categories, such as WD40 repeat domains (WDRs), protein kinases, and armadillo-type fold proteins. WDR-related terms were enriched in both positively and negatively correlated CpG-methylated genes (**Data S4**) and provide a mechanism for CpG methylation to exert large-scale regulation of developmental state. WD40 domains often facilitate protein-protein interaction, especially with other WDR-containing proteins, occupying important hub positions in cellular interaction networks (Stirnimann *et al*., 2010). WDRs are essential for nuclear complexes involved in chromatin remodeling, including two WDRs in the polycomb repressive complex that adds silencing modifications to histones (Margueron *et al*., 2009) and two WDRs in the CLR4 methyltransferase complex (Zhang *et al*., 2008). In the rice internode, a CpG positively correlated WDR-encoding gene, *LOC_Os01g08190*, is an ortholog of LUH (*At2G32700*), which encodes a transcriptional repressor that forms a complex with LEUNING, functioning in regulating flower development, mucilage secretion, pathogen resistance, and cell wall formation (Guerriero *et al*., 2015; Jain & Pandey, 2018). Protein kinase terms are enriched in the lists of genes whose expression is negatively correlated with CpG and positively correlated with CHG. Serine/threonine protein kinases were also enriched in the DM across methylation contexts and in both younger and older segments (**Data S2**). The results are consistent with methylation being an important regulator of kinase transcription throughout internode development and potentially an important regulatory control point for coordination of signaling.

Besides regulators, enzyme-related terms were enriched among methylation-transcript abundance correlations, including cytochrome P450s and other lignin biosynthesis genes, glycoside hydrolase (GHs), and NAD(P)-binding Rossmann fold proteins. Cytochrome P450s are enriched in gene lists of positively and negatively correlated CHG and CHH marks. The CHG positively correlated genes include *LOC_Os10g36848*, which encodes the characterized rice ferulate 5-hydroxylase (aka. CAld5Hs), involved in syringyl monolignol biosynthesis (Takeda *et al*., 2017). Two flavonoid-3-hydroxylases (*LOC_Os09g26980* and *LOC_Os09g26940)* that function in tricin biosynthesis (Lam *et al*., 2019) were present in CHH positive and CHG positive and negative correlations. Several core phenylpropanoid lignin biosynthetic genes, including *LOC_Os06g44620* (Os4CL4, (Gui *et al*., 2011)) and *LOC_Os04g52280* (OsCAD7 / FC1 (Li *et al*., 2009)) were among the negative CHH methylation-transcript correlations. (**Data S5**). Furthermore, glycoside hydrolase (GHs) are enriched in the positively correlated CpG gene set (**Data S4)** and appear but do not pass a stringent FDR cut-off among CHG and CHH positively correlated genes. GHs function in cell wall polysaccharide turnover and remodeling, and varied expression was noted in other plant organ developmental gradient data (Nazipova *et al*., 2022; Xie *et al*., 2022). These observations provide an avenue for DNA methylation to contribute to coordination of cell wall synthesis and remodeling throughout development.

### Methylation machinery transcript abundance and mutants support functions in internode development

Maintenance methylation by CHG methylation processes (**Tabel S1**), and by extension CpG, are implicated in internode development. Greater differential methylation in young compared to old segments in both CHG and CpG methylation (**Figure 3**) aligns with higher expression in early segments of genes involved in CpG and CHG methylation maintenance, particularly CMT3a and MET1-2, respectively, **Table S9; Figure S8**). Thus, CHG and CpG maintenance methylation of hemi-methylated sites in segments containing and near the intercalary meristem appear important for preserving gene expression patterns in newly divided cells, providing continuity of epigenetic memory (Jullien *et al*., 2006; Pikaard & Mittelsten Scheid, 2014). The observation that OsCMT3a loss reduced CHG methylation, activated transposons, and caused pleiotropic developmental abnormalities, including reduced plant height (Cheng *et al*., 2015) supports a role for CHG maintenance in internode cell division (or cellular elongation) defect in this mutant. For CpG maintenance methylation, the rice *met1-2* mutant produced global hypomethylation, abnormal seed development, vivipary, and embryonic lethality (Hu *et al*., 2014; Yamauchi *et al*., 2014), preventing characterization later in development (Liang *et al*., 2022), but the similar expression profiles of MET1-2 and CMT3a support similar roles. In addition, results that CpG methylation within gene bodies is permissive of expression and correlations between CpG methylation and transcript abundance suggest additional roles beyond TE repression, as supported, for example, by CpG methylation reader, MBD707 (Qu *et al*., 2021), in later internode segments (meta-cluster III).

DM, gene expression, genetics, and correlation results also support non-CpG methylation processes functioning in rice internode growth and plant height. *De novo* methylation via the RdDM pathway that primarily controls CHH (Markulin *et al*., 2021) appears to be active as internode development proceeds based on expression in meta-clusters II, III, and IV, including of domains rearranged methyltransferases (DRM2, DRM3), RNA Dependent RNA Polymerase, Dicer-like, Argonaute, and Stabilizer (i.e., WAF1) (**Table S9, Figure S8**, **Figure S10**).

Multiplex CRISPR editing of methyltransferases (DRM2, DRM3, CMT2, CMT3a, CMT3b) demonstrated that progressive removal of non-CpG methylation pathways leads to increasingly severe phenotypes, again including reduced plant growth and height defects (Hu *et al*., 2021).

Small-RNA pathway genetic analysis, which primarily methylate CHH, but also act on in CHG and CpG at some loci (ref) further connect methylation to internode architecture. OsDCL3a regulates hormone pathways, including gibberellin and brassinosteroid signaling, which directly control stem elongation and plant height (Wei *et al*., 2014), and RNA polymerase V is required for proper reproductive development and epigenetic regulation (Zheng *et al*., 2021). Disruption of OsDRM2 produces semi-dwarf plants and reduced tiller number, demonstrating that de novo methylation is required for normal plant height and branching (Moritoh *et al*., 2012). Similarly, mutation of OsDDM1 causes severe early growth defects associated with hypomethylation and genome instability (Tan *et al*., 2016). Consistent with this, distribution of CHH methylation displays a similar profile in early and later segments, coinciding with increased expression of OsCMT2 and OsDRM2, which maintain CHH methylation.

Finally, loss of the demethylase ROS1a alters plant architecture in a different way, with reports of increased tillering, reduced panicle length, but no affect on plant height per se, though internode lengths were not measured (Irshad *et al*., 2022). This suggests that balanced methylation and demethylation are required to maintain normal plant height and branching patterns. The increase of ROS1a and ROS1d later in internode development (meta cluster III and IV) are consistent with active demethylation as part of internode expression regulation.

## Conclusion

This study supports a role for DNA methylation in regulating gene expression across the pseudo-time course of the rice internode. Our model (**Figure S11**) summarizes distinct methylation dynamics, showing a shift from CpG and CHG dominated patterns in younger segments to similar prevalence with CHH methylation in older internode segments. This suggests that the need for continued CpG- and CHG-mediated gene regulation, including of TEs, diminishes with development. Both in TE and non-TE genes, CpG methylation in coding regions and CHG/CHH methylation in upstream regions is associated with permissive gene expression in our dataset. The data indicate that single marks within the context of CpG_CHG and triple marked non-TE genes are typically the main drivers of methylation-expression correlations. revealing associations between methylation and particular, non-random gene families. Across all three methylation contexts (CpG, CHG, and CHH), serine/threonine protein kinases are consistently enriched among differentially methylated genes, highlighting kinase- and WDR-mediated signaling as a central partner of DNA methylation in the coordinated regulation of rice internode development. Further, correlations between methylation and expression of lignin biosynthesis genes and cell wall-related glycoside hydrolases support a model that methylation functions to alter cell wall development. These observations offer avenues for adjusting expression of groups of genes for biotechnological purposes such as altering the period of elongation to increase or decrease internode length, or coordinately alter secondary cell wall development.

## Authors’ Contributions

L.B. and F.L. designed the study.

F.L. collected samples and prepared nucleic acids.

A.L., V.S., V.N, C.D., Y.Y generated data

M.M., J.D., N.G., S.F., L.B., N.N.C conceived and executed the data analysis.

D.X., L.S., S.L assisted with the computational processing of data.

S.F., J.D., L.B. provided guidance in the interpretation of results.

N.N.C., M.M., and L.B. drafted the manuscript.

M.M., N.N.C, N.G, J.D., L.B., D.X edited the manuscript.

All authors read and approved the manuscript.

## Acknowledgments

We thank Ms. Jillian Vaught for assistance with sample preparation, Dr. J.B. Jewell, Dr. R. Panahabadi, Mr. Jordon Tolley, and Dr. Christine Queitsch for insightful discussion.

## Data Availability Statement

Data supporting the findings of this study are provided within the figures and supplementary materials of this article. Raw transcriptomic and bisulfite sequencing datasets have been deposited in the NCBI Sequence Read Archive (SRA), with accession numbers listed in the supplementary materials (**Data S6**).

## Supporting Figures & Tables

**Tabel S1:** Loss-of-function mutants in rice DNA methylation pathways and their effects on plant height, growth and development.

| Gene / Pathway | Primary Role | Methylation Context | Effect on Plant Height / Architecture | Reference |
| --- | --- | --- | --- | --- |
| **MET1b** | **Writer:** Maintenance methyltransferase - | CpG | *met1b* homozygous mutants showed abnormal seed development associated with viviparous germination or early embryonic lethality, preventing normal growth. | Yamauchi et al., 2014 |
| **Major MET1 CG methylase** | **Writer:** CG maintenance methylation | CpG | Homozygous *met1* seedlings grew only to approximately 3 cm, developed sporadic/deformed tiny roots, then ceased growth and underwent necrotic death within 2 weeks. | Hu et al., 2014 |
| **CMT3a** | **Writer:** Chromomethylase | CHG | Pleiotropic developmental abnormalities, including early heading, decreased plant height, and reduced fertility. | Cheng et al., 2015 |
| **DRM2, DRM3, CMT2, CMT3a, and CMT3b** | **Writer:** Redundant non-CpG methyltransferases | Primarily CHG and CHH | Reduced plant height and growth suppression, with stronger phenotypes in higher-order mutants carrying combined knockouts of multiple non-CG methyltransferase genes. | Hu et al., 2021 |
| **DCL3a** | **RdDM pathway:** siRNA production | CHH (RdDM-associated) | RNAi knockdown lines showed decreased plant height/dwarf phenotype, altered flag leaf inclination, reduced panicle length, and changes in the number of main panicle branches. | Wei et al., 2014 |
| **RNA Polymerase V** | **RdDM pathway** | CHH (locus-specific) | *osnrpe1* (largest subunit of RNA pol V) mutants showed reduced plant height and failed to produce seeds. | Zheng et al., 2021 |
| **DRM2** | **Writer:** *De novo* methyltransferase - | CHH (also CpG and CHG at some loci) | Semi-dwarf plants, reduced tiller number, delayed heading, abnormal panicles | Moritoh et al., 2012 |
| **DDM1** | **Reader:** Chromatin remodeling enabling methylation | Primarily CHG and CHH (also CpG in heterochromatin) | osddm1a/1b mutants showed severely reduced plant height | Tan et al., 2016 |
| **ROS1a** | **Eraser:** DNA demethylase | Removes CpG, CHG, and CHH | Tiller number was increased by 44% and seed fertility was significantly reduced by 23%. | Irshad et al., 2022 |

**Figure S1:**
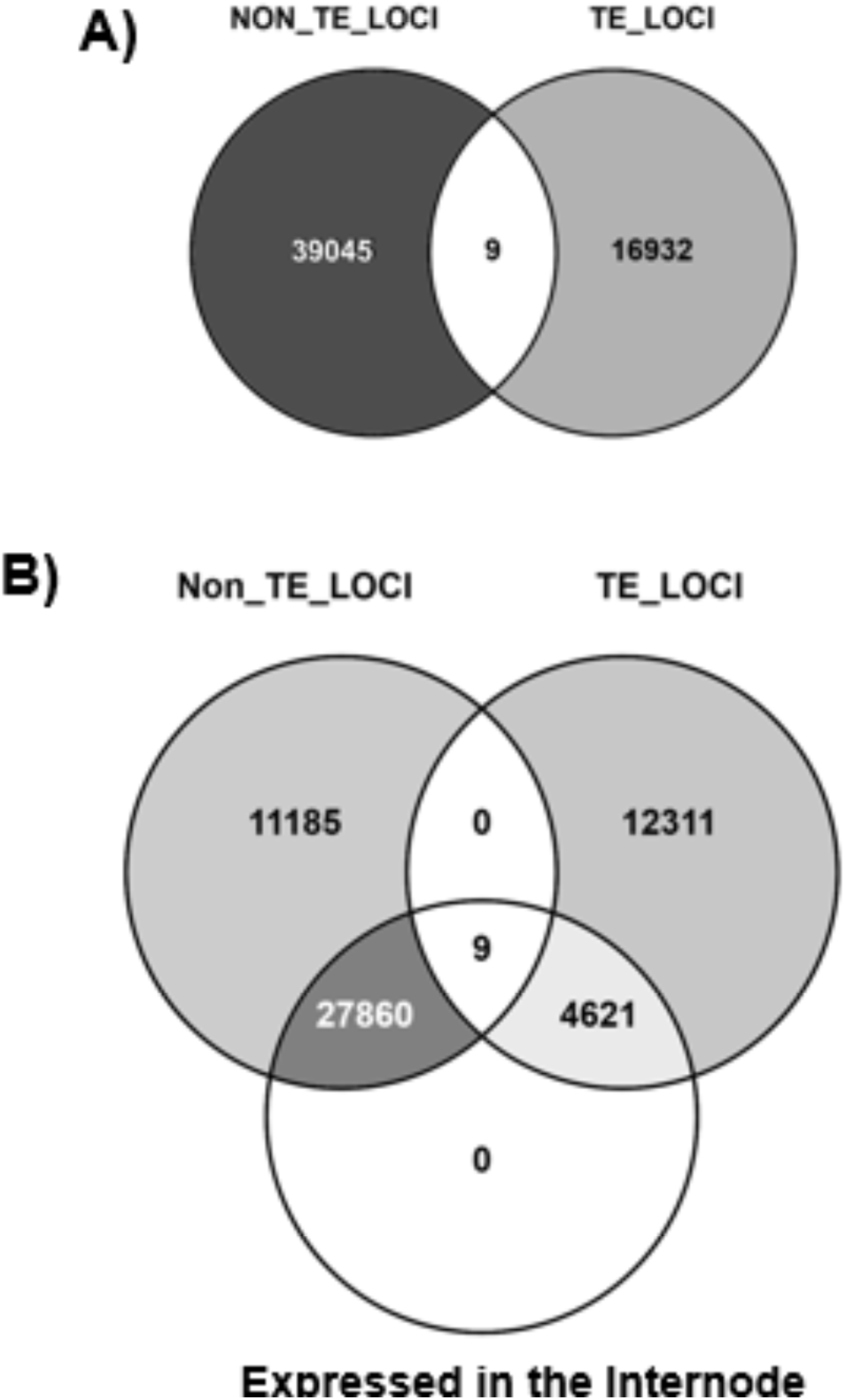
Comparison of transposable element (TE)-associated and non-TE-associated loci with expressed genes in our dataset. **(A)** Venn diagram showing the overlap between TE-associated loci (TE_LOCI) and non-TE-associated loci (NON_TE_LOCI) in the rice genome, as obtained from the Rice Genome Annotation Project (Rice UGA, https://rice.uga.edu/). A total of 9 loci are shared between TE and non-TE categories. **(B)** Venn diagram comparing the expressed loci in our dataset (Expressed in the internode) with TE_LOCI and NON_TE_LOCI categories.

**Figure S2.**
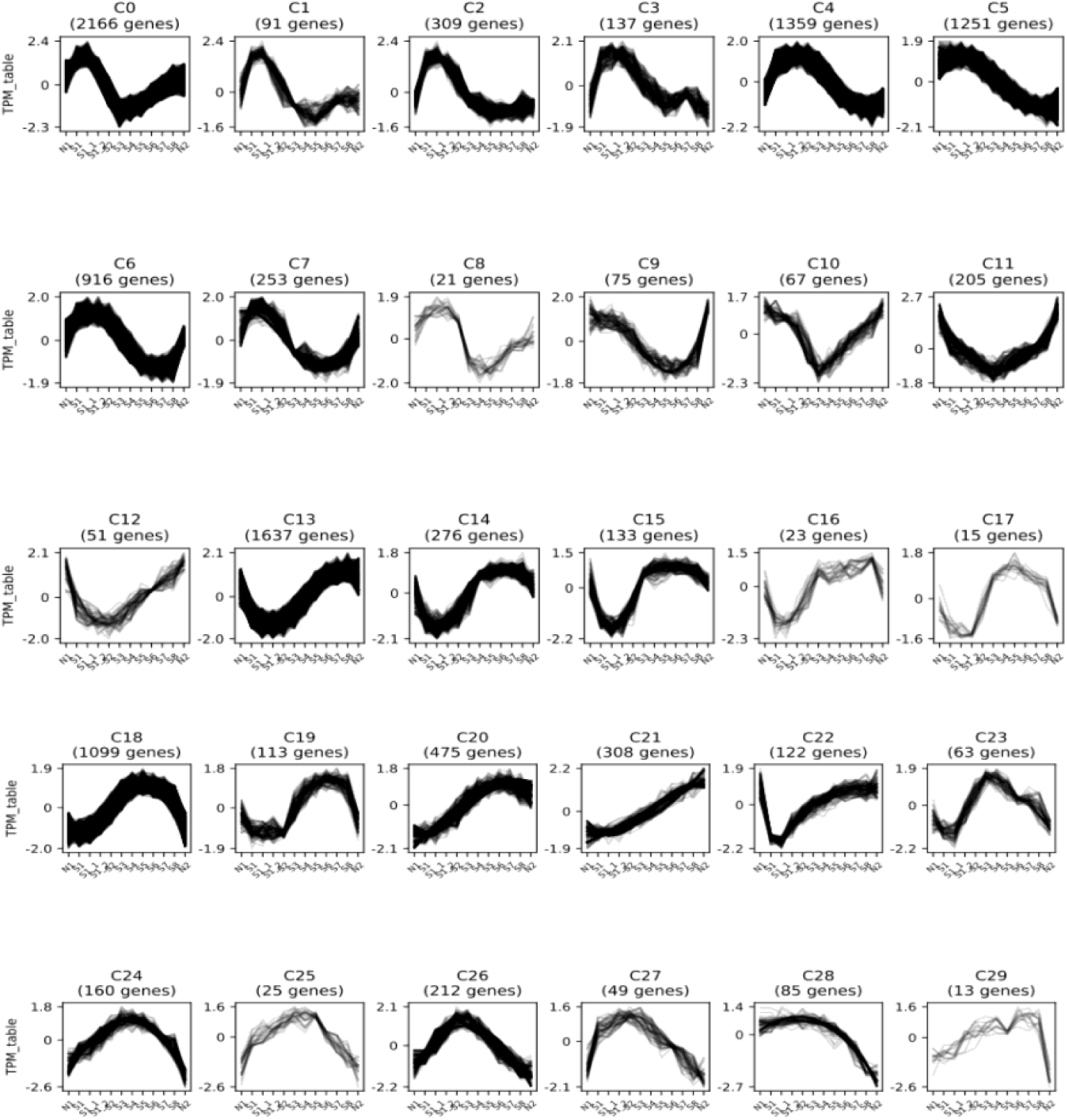
CLUST cluster profiles. Raw TPM values for all genes were log2 transformed and z-score quantile normalized. These normalized values were used to cluster genes into co-expression clusters using the CLUST software tool resulting in 30 gene co-expression clusters.

**Figure S3:**
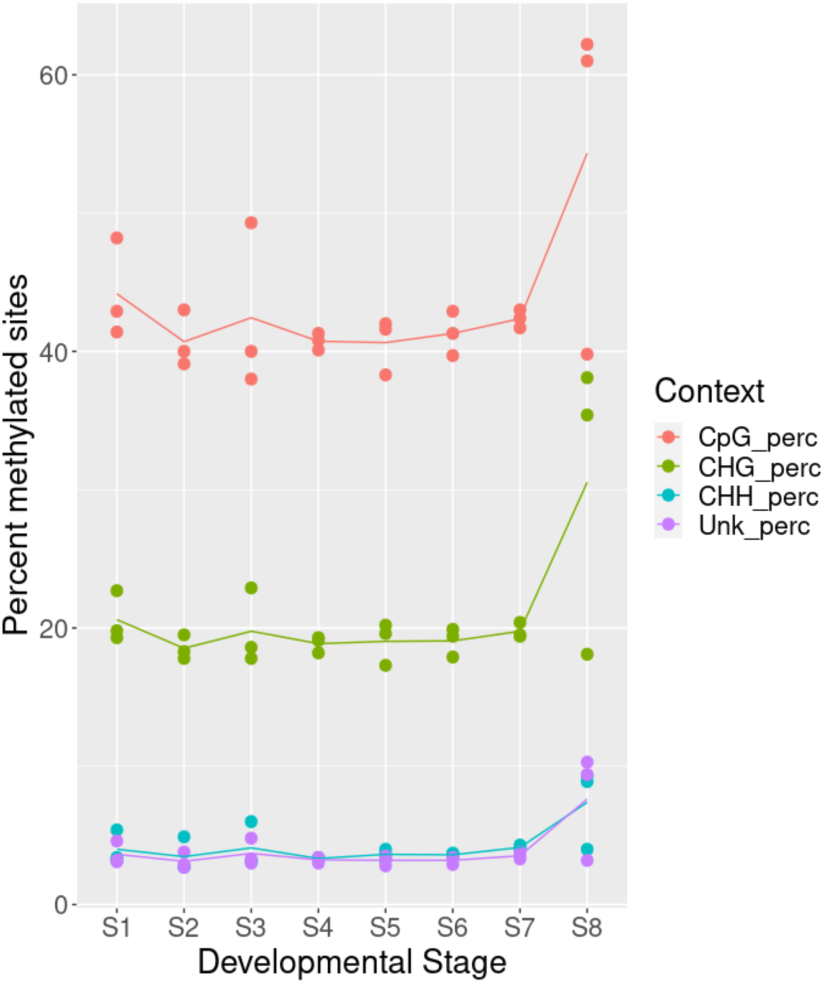
Methylation profiles across developmental stages. The graph represents the percentage of methylated sites across different methylation contexts (CpG, CHG, CHH, and Unknown) for rice developmental stages S1–S8. CpG methylation remained relatively stable (∼40%) across stages S1–S7. CHG methylation averaged ∼20%, showing slight variation across stages, while CHH methylation remained low (∼4%) throughout the stages. Unknown contexts displayed minimal methylation. Two S8 samples exhibited anomalously higher methylation percentages across all contexts, suggesting experimental errors; these samples were excluded from downstream analyses. Points represent biological replicates, and lines indicate context-specific trends over developmental stages.

**Table S2:**
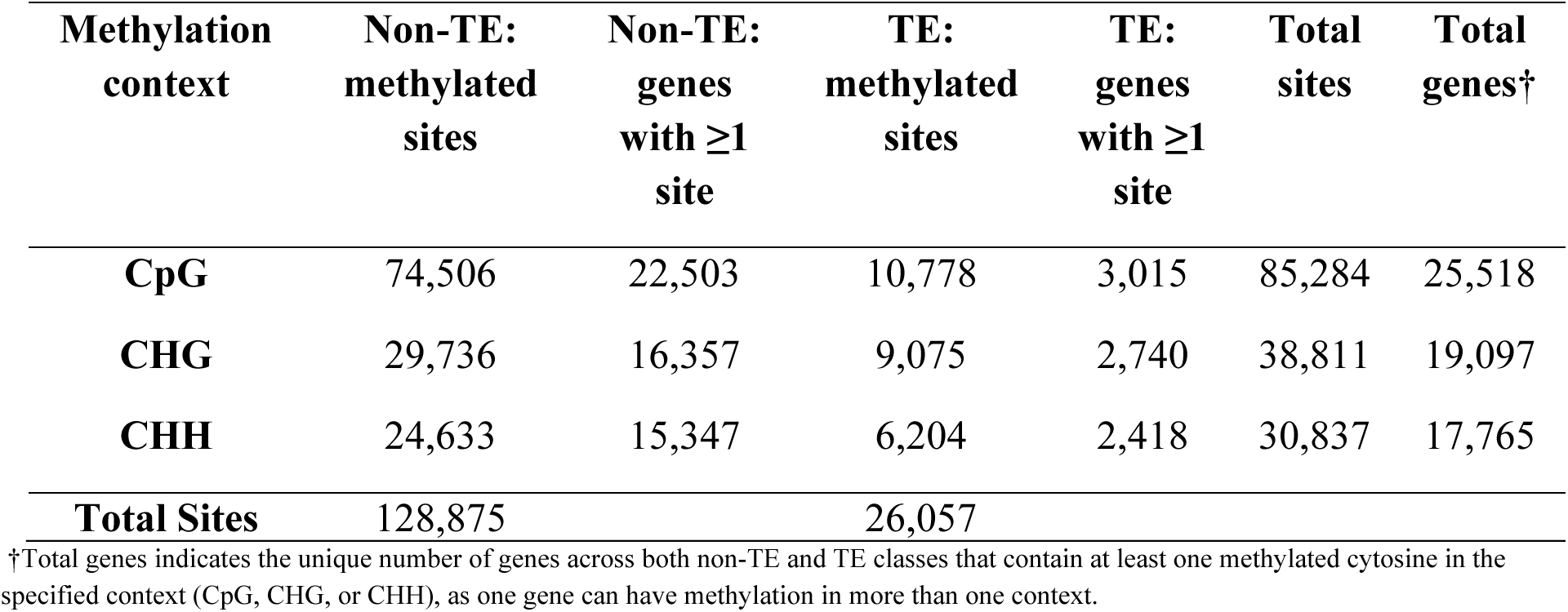
Summary of Methylation Events in non-TE and TE Genes.

**Figure S4:**
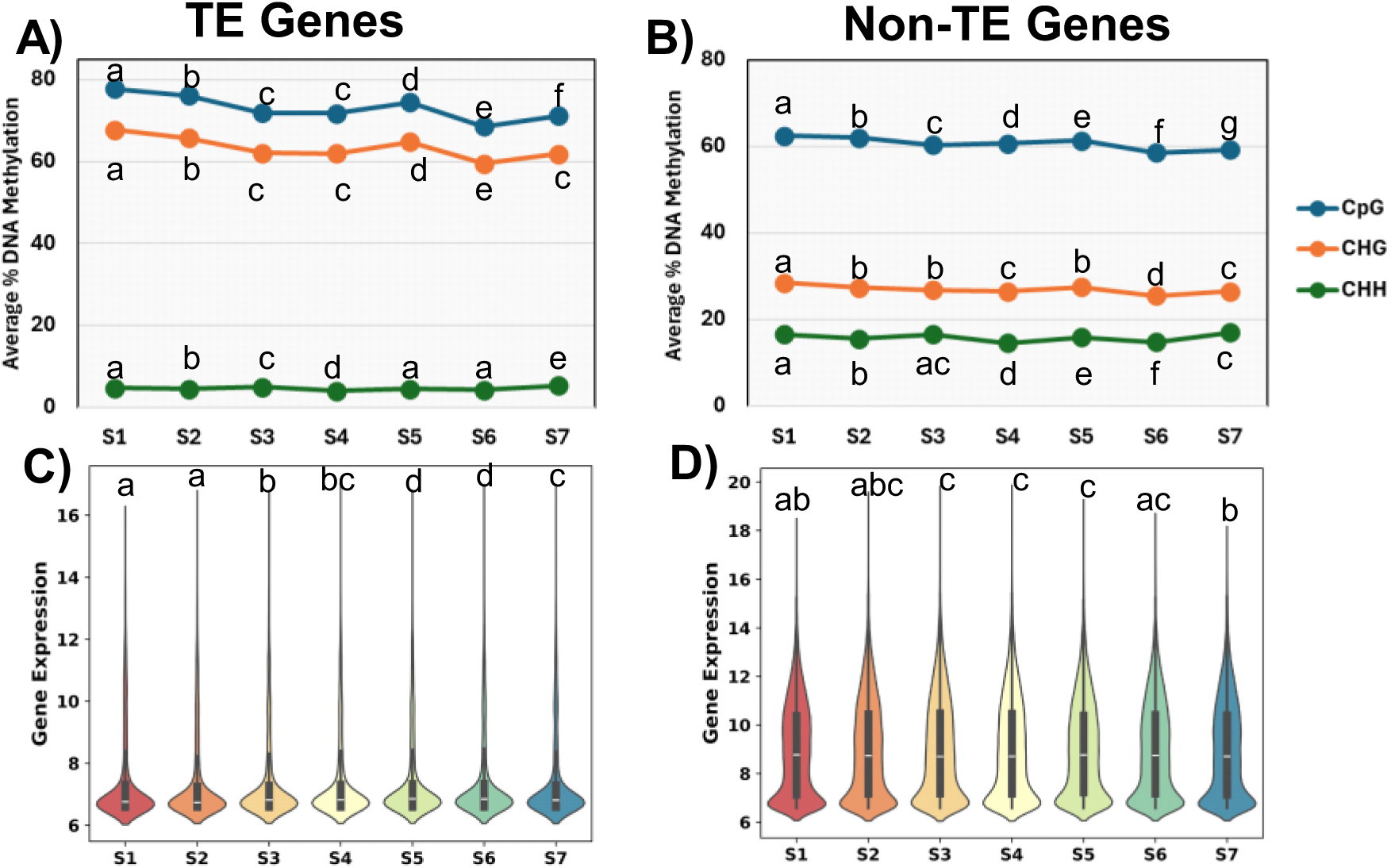
Distinct DNA methylation patterns and gene expression across internode segments for TE and Non-TE Genes. **(A)** and **(B)** DNA methylation levels across internode segments. The final output of the DNA methylation analysis (see Methods) consisted of a matrix for each methylation context, containing the percentage regions as rows and segments as columns, storing the percentage (0%-100%) of methylated base pairs for each combination. Panels show the mean methylation percentages for each internode segment (S1–S7) in the CpG, CHG, and CHH contexts. **(C)** and **(D)** Distribution of gene expression of the methylated genes across the segments. The raw counts were variance-stabilized transformation (VST) for gene expression, with no or low expression corresponding to a value of approximately ∼6.5. Group differences were assessed using a Kruskal-Wallis test followed, where significant, by Dunn’s post-hoc multiple-comparisons test; compact letter displays indicate statistically distinct groups (groups sharing a letter are not significantly different).

**Figure S5:**
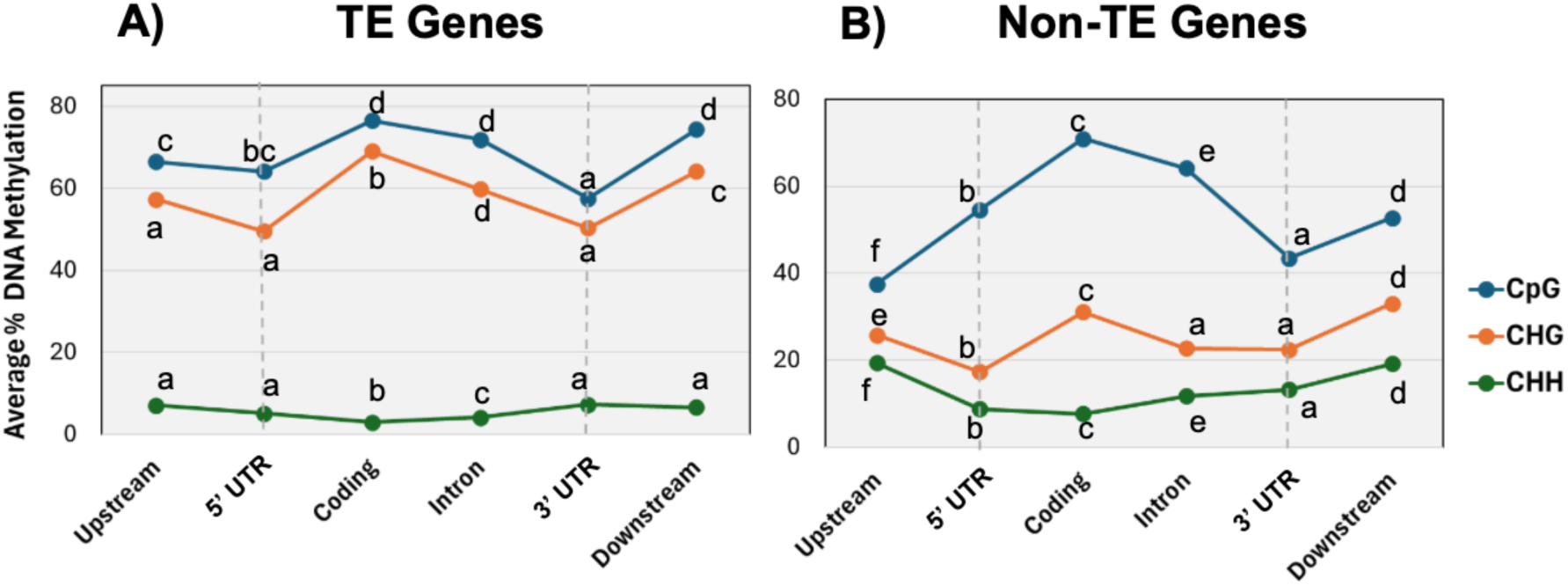
DNA methylation profiles across gene features. The average percentage of DNA methylation in CpG, CHG, and CHH contexts is shown for different genomic regions, including upstream, 5’ untranslated region (5’UTR), coding regions, introns, 3’ untranslated region (3’UTR), and downstream sequences in **(A)** TE-genes and **(B)** non-TE genes. The patterns of methylation differ between A and B, reflecting their divergent regulatory mechanisms. Group differences were assessed using a Kruskal-Wallis test followed, where significant, by Dunn’s post-hoc multiple-comparisons test; compact letter displays indicate statistically distinct groups (groups sharing a letter are not significantly different).

**Table S3.**
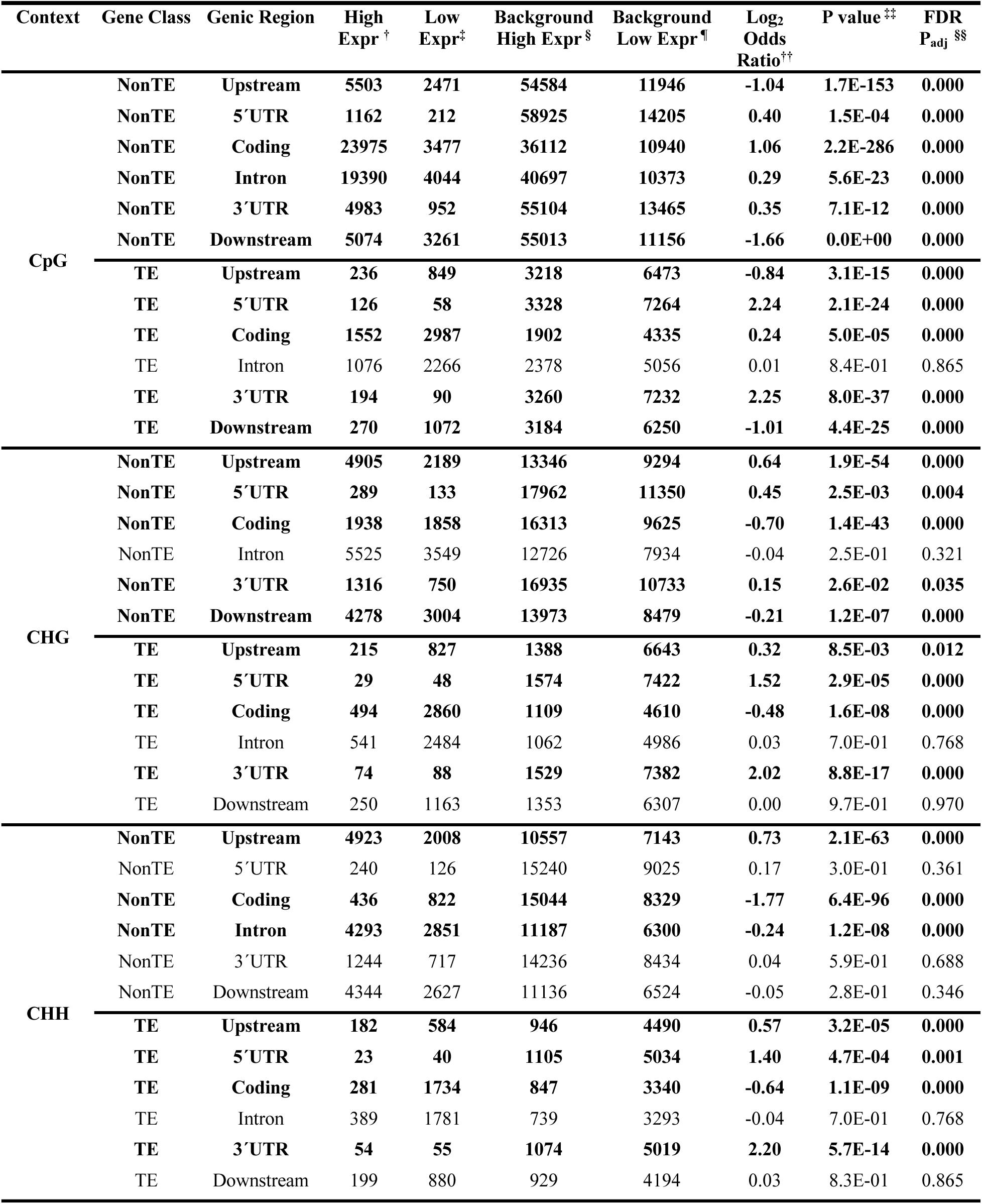
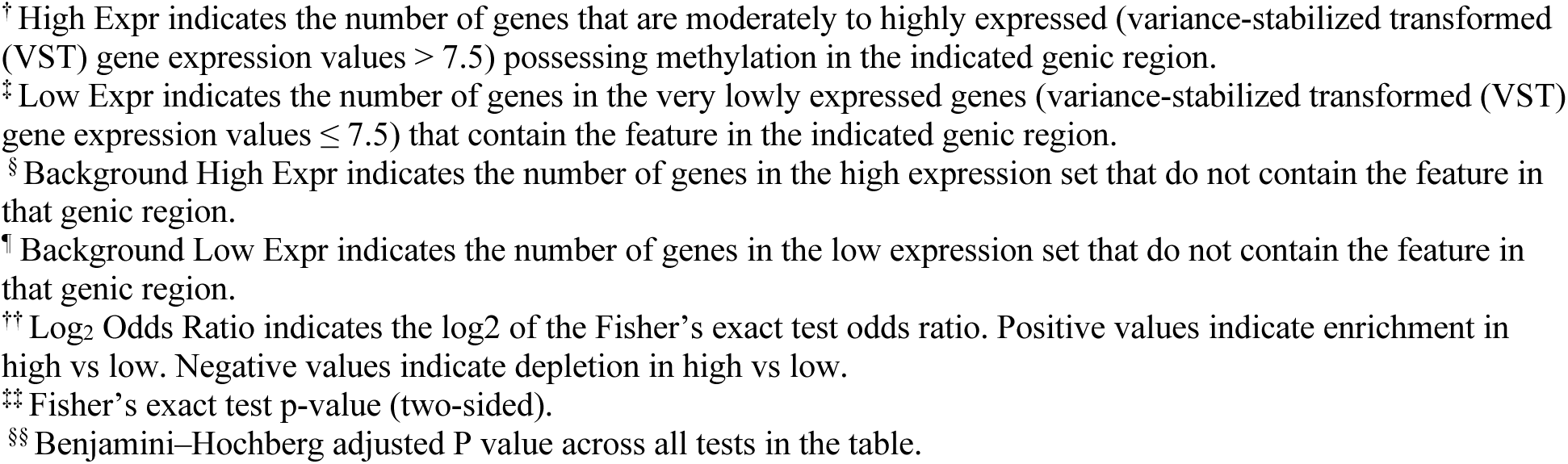
Associations between DNA methylation context and gene expression in different genic regions to support **Figure 4**. Rows shown in bold indicate associations that passed the FDR significance threshold (P_adj_ ≤ 0.05). Among expressed and methylated non-TE genes (N = 23,643), 70% were classified as moderately to highly expressed, defined as mean expression > 7.5 VST across S1–S7, whereas 30% were classified as lowly expressed, defined as mean expression ≤ 7.5 VST. In contrast, among expressed and methylated TE-associated genes (N = 3,309), 22.5% were moderately to highly expressed and 77.5% were lowly expressed using the same expression threshold.

| Context | Gene Class | Genic Region | High Expr<sup>†</sup> | Low Expr<sup>‡</sup> | Background High Expr<sup>§</sup> | Background Low Expr<sup>¶</sup> | Log<sub>2</sub> Odds Ratio<sup>††</sup> | P value<sup>‡‡</sup> | FDR P<sub>adj</sub><sup>§§</sup> |
| --- | --- | --- | --- | --- | --- | --- | --- | --- | --- |
| CpG | NonTE | Upstream | 5503 | 2471 | 54584 | 11946 | -1.04 | 1.7E-153 | 0.000 |
|  | NonTE | 5'UTR | 1162 | 212 | 58925 | 14205 | 0.40 | 1.5E-04 | 0.000 |
|  | NonTE | Coding | 23975 | 3477 | 36112 | 10940 | 1.06 | 2.2E-286 | 0.000 |
|  | NonTE | Intron | 19390 | 4044 | 40697 | 10373 | 0.29 | 5.6E-23 | 0.000 |
|  | NonTE | 3'UTR | 4983 | 952 | 55104 | 13465 | 0.35 | 7.1E-12 | 0.000 |
|  | NonTE | Downstream | 5074 | 3261 | 55013 | 11156 | -1.66 | 0.0E+00 | 0.000 |
|  | TE | Upstream | 236 | 849 | 3218 | 6473 | -0.84 | 3.1E-15 | 0.000 |
|  | TE | 5'UTR | 126 | 58 | 3328 | 7264 | 2.24 | 2.1E-24 | 0.000 |
|  | TE | Coding | 1552 | 2987 | 1902 | 4335 | 0.24 | 5.0E-05 | 0.000 |
|  | TE | Intron | 1076 | 2266 | 2378 | 5056 | 0.01 | 8.4E-01 | 0.865 |
|  | TE | 3'UTR | 194 | 90 | 3260 | 7232 | 2.25 | 8.0E-37 | 0.000 |
|  | TE | Downstream | 270 | 1072 | 3184 | 6250 | -1.01 | 4.4E-25 | 0.000 |
| CHG | NonTE | Upstream | 4905 | 2189 | 13346 | 9294 | 0.64 | 1.9E-54 | 0.000 |
|  | NonTE | 5'UTR | 289 | 133 | 17962 | 11350 | 0.45 | 2.5E-03 | 0.004 |
|  | NonTE | Coding | 1938 | 1858 | 16313 | 9625 | -0.70 | 1.4E-43 | 0.000 |
|  | NonTE | Intron | 5525 | 3549 | 12726 | 7934 | -0.04 | 2.5E-01 | 0.321 |
|  | NonTE | 3'UTR | 1316 | 750 | 16935 | 10733 | 0.15 | 2.6E-02 | 0.035 |
|  | NonTE | Downstream | 4278 | 3004 | 13973 | 8479 | -0.21 | 1.2E-07 | 0.000 |
|  | TE | Upstream | 215 | 827 | 1388 | 6643 | 0.32 | 8.5E-03 | 0.012 |
|  | TE | 5'UTR | 29 | 48 | 1574 | 7422 | 1.52 | 2.9E-05 | 0.000 |
|  | TE | Coding | 494 | 2860 | 1109 | 4610 | -0.48 | 1.6E-08 | 0.000 |
|  | TE | Intron | 541 | 2484 | 1062 | 4986 | 0.03 | 7.0E-01 | 0.768 |
|  | TE | 3'UTR | 74 | 88 | 1529 | 7382 | 2.02 | 8.8E-17 | 0.000 |
|  | TE | Downstream | 250 | 1163 | 1353 | 6307 | 0.00 | 9.7E-01 | 0.970 |
| CHH | NonTE | Upstream | 4923 | 2008 | 10557 | 7143 | 0.73 | 2.1E-63 | 0.000 |
|  | NonTE | 5'UTR | 240 | 126 | 15240 | 9025 | 0.17 | 3.0E-01 | 0.361 |
|  | NonTE | Coding | 436 | 822 | 15044 | 8329 | -1.77 | 6.4E-96 | 0.000 |
|  | NonTE | Intron | 4293 | 2851 | 11187 | 6300 | -0.24 | 1.2E-08 | 0.000 |
|  | NonTE | 3'UTR | 1244 | 717 | 14236 | 8434 | 0.04 | 5.9E-01 | 0.688 |
|  | NonTE | Downstream | 4344 | 2627 | 11136 | 6524 | -0.05 | 2.8E-01 | 0.346 |
|  | TE | Upstream | 182 | 584 | 946 | 4490 | 0.57 | 3.2E-05 | 0.000 |
|  | TE | 5'UTR | 23 | 40 | 1105 | 5034 | 1.40 | 4.7E-04 | 0.001 |
|  | TE | Coding | 281 | 1734 | 847 | 3340 | -0.64 | 1.1E-09 | 0.000 |
|  | TE | Intron | 389 | 1781 | 739 | 3293 | -0.04 | 7.0E-01 | 0.768 |
|  | TE | 3'UTR | 54 | 55 | 1074 | 5019 | 2.20 | 5.7E-14 | 0.000 |
|  | TE | Downstream | 199 | 880 | 929 | 4194 | 0.03 | 8.3E-01 | 0.865 |

<sup>†</sup> High Expr indicates the number of genes that are moderately to highly expressed (variance-stabilized transformed (VST) gene expression values $> 7.5$ ) possessing methylation in the indicated genic region.
<sup>‡</sup> Low Expr indicates the number of genes in the very lowly expressed genes (variance-stabilized transformed (VST) gene expression values $\leq 7.5$ ) that contain the feature in the indicated genic region.
<sup>§</sup> Background High Expr indicates the number of genes in the high expression set that do not contain the feature in that genic region.
<sup>¶</sup> Background Low Expr indicates the number of genes in the low expression set that do not contain the feature in that genic region.
<sup>††</sup> Log<sub>2</sub> Odds Ratio indicates the log<sub>2</sub> of the Fisher's exact test odds ratio. Positive values indicate enrichment in high vs low. Negative values indicate depletion in high vs low.
<sup>‡‡</sup> Fisher's exact test p-value (two-sided).
<sup>§§</sup> Benjamini–Hochberg adjusted P value across all tests in the table.

**Figure S6:**
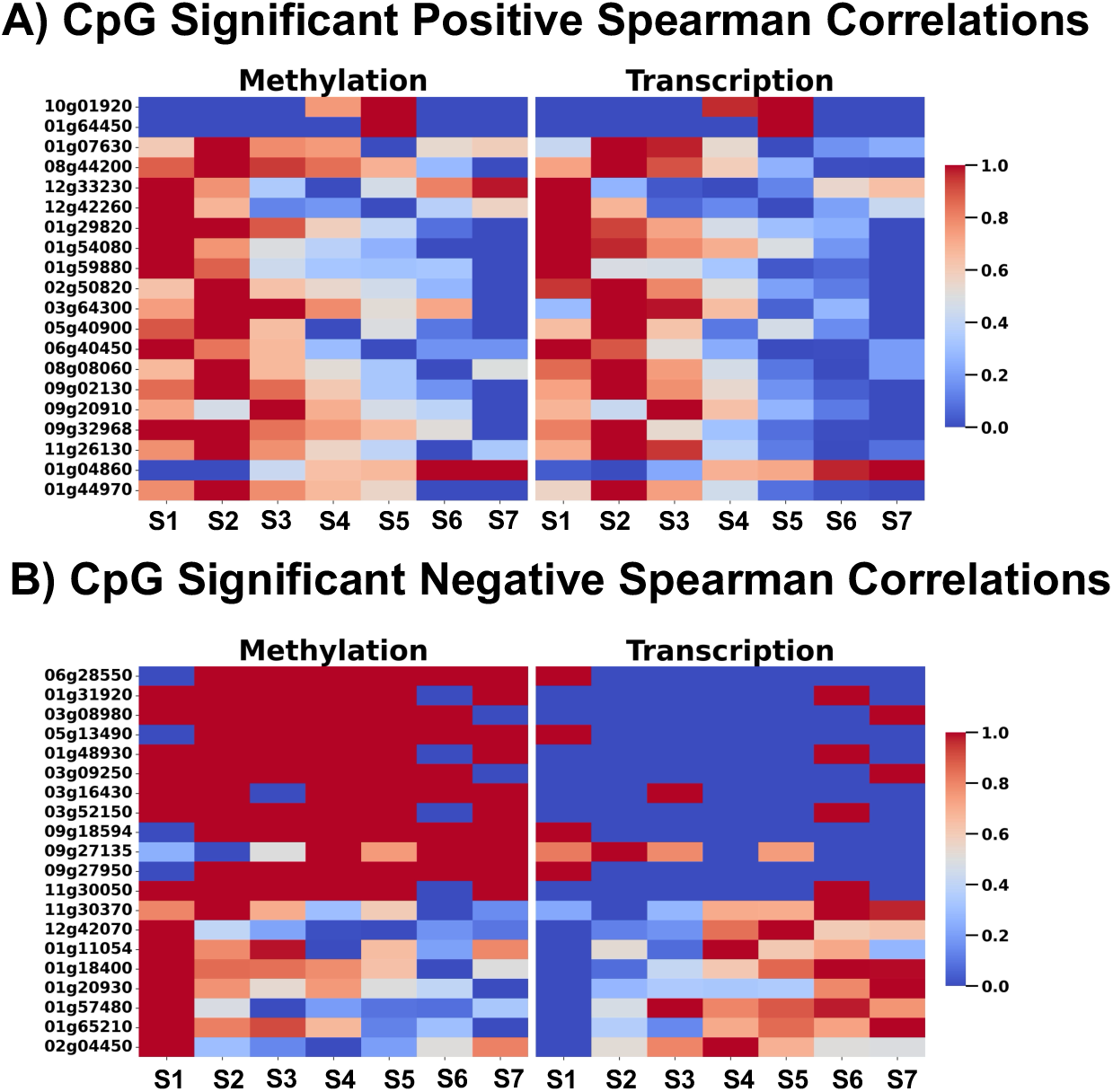
Strongest CpG methylation-transcription correlations among non-TE genes across rice internode development. Heatmaps visualize the 20 strongest positive **(A)** and negative **(B)** correlations between CpG methylation frequency and transcript abundance of non-TE genes across internode segments (S1–S7), with values scaled using min–max normalization. The data used to generate this heatmap is in **Data S3**.

**Table S4.**
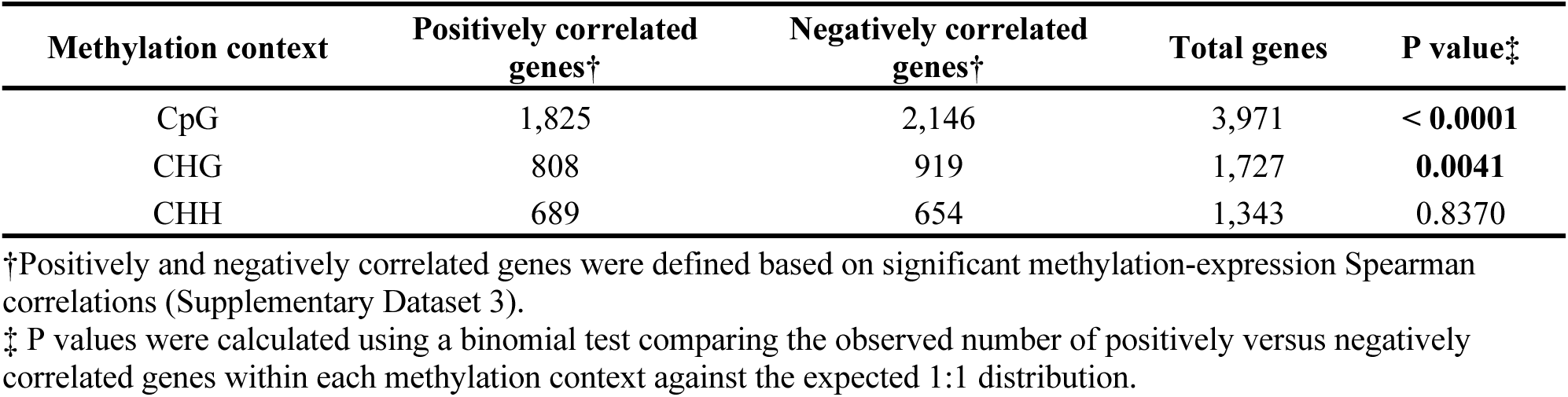
Counts of genes showing significant positive or negative methylation-expression correlations for each methylation context.

**Table S5.** Regional DNA methylation differences between positively and negatively correlated genes by methylation context.

| Methylation context | Genomic region | Positive mean methylation (%)<sup>†</sup> | Negative mean methylation (%)<sup>†</sup> | Positive median methylation (%)<sup>†</sup> | Negative median methylation (%)<sup>†</sup> | Mean difference: Positive – Negative (%) | Median difference: Positive – Negative (%) | Mann-Whitney U statistic<sup>‡</sup> | Raw P value<sup>‡</sup> | BH-adjusted P value<sup>§</sup> |
| --- | --- | --- | --- | --- | --- | --- | --- | --- | --- | --- |
| CpG | Upstream | 34.62 | 38.03 | 28.23 | 29.77 | -3.40 | -1.55 | 33040 | 1.83E-01 | 0.548 |
|  | 5' UTR | 55.31 | 54.94 | 61.69 | 63.83 | 0.37 | -2.14 | 967.5 | 9.34E-01 | 0.968 |
|  | Coding | 66.63 | 63.83 | 79.37 | 75.52 | 2.80 | 3.85 | 293820.5 | 2.51E-02 | 0.262 |
|  | Intron | 58.38 | 60.35 | 63.09 | 65.90 | -1.97 | -2.81 | 249431.5 | 3.20E-01 | 0.720 |
|  | 3' UTR | 45.11 | 42.68 | 43.93 | 36.69 | 2.43 | 7.24 | 20557 | 4.21E-01 | 0.843 |
|  | Downstream | 47.49 | 53.10 | 45.83 | 54.05 | -5.61 | -8.21 | 31561.5 | 4.37E-02 | 0.262 |
| CHG | Upstream | 26.43 | 26.02 | 18.80 | 17.65 | 0.41 | 1.15 | 25781.5 | 8.54E-01 | 0.968 |
|  | 5' UTR | 13.98 | 14.72 | 4.95 | 2.87 | -0.73 | 2.08 | 63.5 | 8.70E-01 | 0.968 |
|  | Coding | 25.52 | 30.43 | 6.43 | 16.33 | -4.90 | -9.90 | 4705 | 3.04E-02 | 0.262 |
|  | Intron | 21.75 | 23.55 | 12.21 | 14.51 | -1.80 | -2.30 | 37693 | 4.74E-01 | 0.853 |
|  | 3' UTR | 22.20 | 26.27 | 8.08 | 17.32 | -4.07 | -9.24 | 1796 | 1.52E-01 | 0.547 |
|  | Downstream | 29.90 | 34.46 | 22.05 | 27.43 | -4.56 | -5.38 | 21542 | 5.95E-02 | 0.268 |
| CHH | Upstream | 19.54 | 19.21 | 15.78 | 14.48 | 0.33 | 1.30 | 18103.5 | 9.68E-01 | 0.968 |
|  | 5' UTR | 7.65 | 6.56 | 2.10 | 3.30 | 1.10 | -1.20 | 50 | 7.14E-01 | 0.968 |
|  | Coding | 5.99 | 6.85 | 1.18 | 1.43 | -0.86 | -0.25 | 349.5 | 6.43E-01 | 0.968 |
|  | Intron | 10.10 | 11.73 | 5.76 | 7.90 | -1.64 | -2.14 | 20064 | 2.67E-01 | 0.686 |
|  | 3' UTR | 12.70 | 11.22 | 8.36 | 8.90 | 1.49 | -0.55 | 1433 | 7.70E-01 | 0.968 |
|  | Downstream | 17.90 | 18.10 | 14.45 | 12.55 | -0.20 | 1.90 | 19990.5 | 7.97E-01 | 0.968 |

<sup>†</sup>Positive and negative groups refer to genes with significant positive or negative methylation–expression Spearman correlations, respectively. Methylation values represent average regional DNA methylation percentages for each methylation context and genic region.
<sup>‡</sup>Positive and negative methylation distributions were compared within each methylation context and genic region using a two-sided Mann-Whitney U test. Raw P values are shown.
<sup>§</sup>P values were adjusted across all 18 comparisons using the Benjamini-Hochberg false discovery rate correction. No comparisons passed the FDR significance threshold of $P_{adj} \leq 0.05$ .

**Figure S7:**
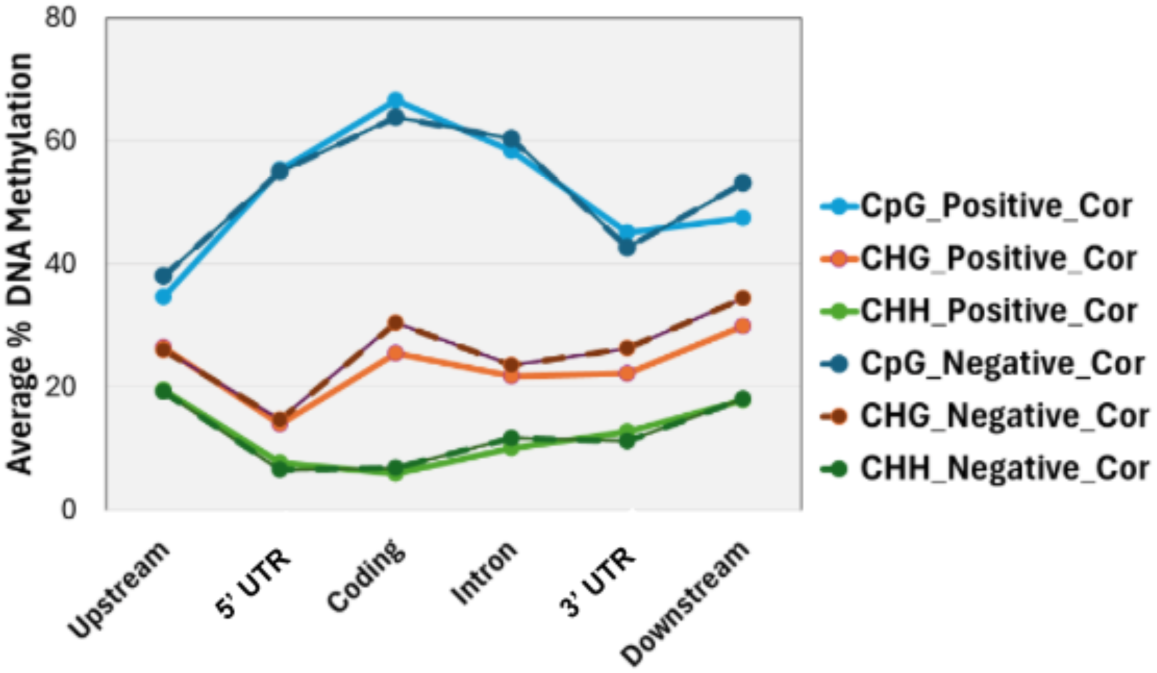
DNA methylation profiles across gene features for significantly correlated genes. DNA methylation patterns across gene features of positively and negatively correlated non-TE genes. The average percentage of DNA methylation in CpG, CHG, and CHH contexts is shown across various genomic regions, including upstream, 5’UTR, coding, introns, 3’UTR, and downstream regions. Positively correlated genes are represented by solid lighter lines, and negatively correlated genes by darker dashed lines. This graph emphasizes that there are no differences in the methylation patterns of positively versus negatively correlated genes across the genic features.

**Tabel S6.** Frequency of significant non-TE gene expression vs. methylation correlation data supporting Figure 5B. This table summarizes significant DNA methylation-expression correlations for non-TE elements (significantly correlated subset N = 6,858; background set of methylated non-TE elements N = 56,789). Columns indicate the methylation-mark class carried by each locus: single marks (CpG single, CHG single, CHH single), double marks (CpG_CHG, CpG_CHH, CHG_CHH), or triple marks (CpG_CHG_CHH). Rows indicate combined correlation categories (methylation-mark class × correlation sign), where POS denotes a positive methylation-expression association and NEG denotes a negative association. Starting Counts (Left Most Panel): number of loci in each methylation-mark class in the full background set (N = 56,789). Body cells: number of loci in the significantly correlated subset (N = 6,858) that fall into the specified row (correlation category) and column (methylation-mark class). Total (Middle Panel): column sums (total significantly correlated loci per methylation-mark class). Total (Right Most Panel): row sums (total significantly correlated loci per correlation category). Cells shaded light blue indicate methylation-mark classes that are significantly enriched in the Middle Panel (i.e., over-represented among significantly correlated loci relative to their background frequency). Cells shaded red indicate correlation categories that are significantly enriched in the Right Most Panel (i.e., over-represented relative to expectation based on the global methylation distribution). Enrichment/depletion was evaluated by a hypergeometric test (P < 0.001) using the Starting Counts as the background. Black cells indicate not applicable/structural zeros (a locus cannot contribute to incompatible categories, e.g., a CpG single locus cannot be counted under CHG single, CHH single, or multi-mark columns)

| Starting Counts (Left Most Panel) | 26118 | 1951 | 2239 | 6105 | 2687 | 3097 | 14592 | 56789 |
| --- | --- | --- | --- | --- | --- | --- | --- | --- |
| Significant Correlated Categories | CpG Single | CHG Single | CHH Single | CpG_CHG | CpG_CHH | CHG_CHH | Triple | Total (Right Most Panel) |
| CpG Single_POS | 2316 |  |  | 488 | 170 |  | 898 | 3872 |
| CpG Single_NEG |  |  |  |  |  |  |  |  |
| CHG Single_POS |  | 124 |  | 337 |  | 162 | 706 | 1329 |
| CHG Single_NEG |  |  |  |  |  |  |  |  |
| CHH Single_POS |  |  | 129 |  | 125 | 173 | 672 | 1099 |
| CHH Single_NEG |  |  |  |  |  |  |  |  |
| CpG_CHG_Double_POS |  |  |  | 75 |  |  | 206 | 281 |
| CpG_CHG_Double_NEG |  |  |  |  |  |  |  |  |
| CpG_CHH_Double_POS |  |  |  |  | 20 |  | 88 | 108 |
| CpG_CHH_Double_NEG |  |  |  |  |  |  |  |  |
| CHG_CHH_Double_POS |  |  |  |  |  | 27 | 100 | 127 |
| CHG_CHH_Double_NEG |  |  |  |  |  |  |  |  |
| CpG_CHG_CHH_Triple_POS |  |  |  |  |  |  | 42 | 42 |
| CpG_CHG_CHH_Triple_NEG |  |  |  |  |  |  |  |  |
| Total (Middle Panel) | 2316 | 124 | 129 | 900 | 315 | 362 | 2712 | 6858 |

**Table S7.**
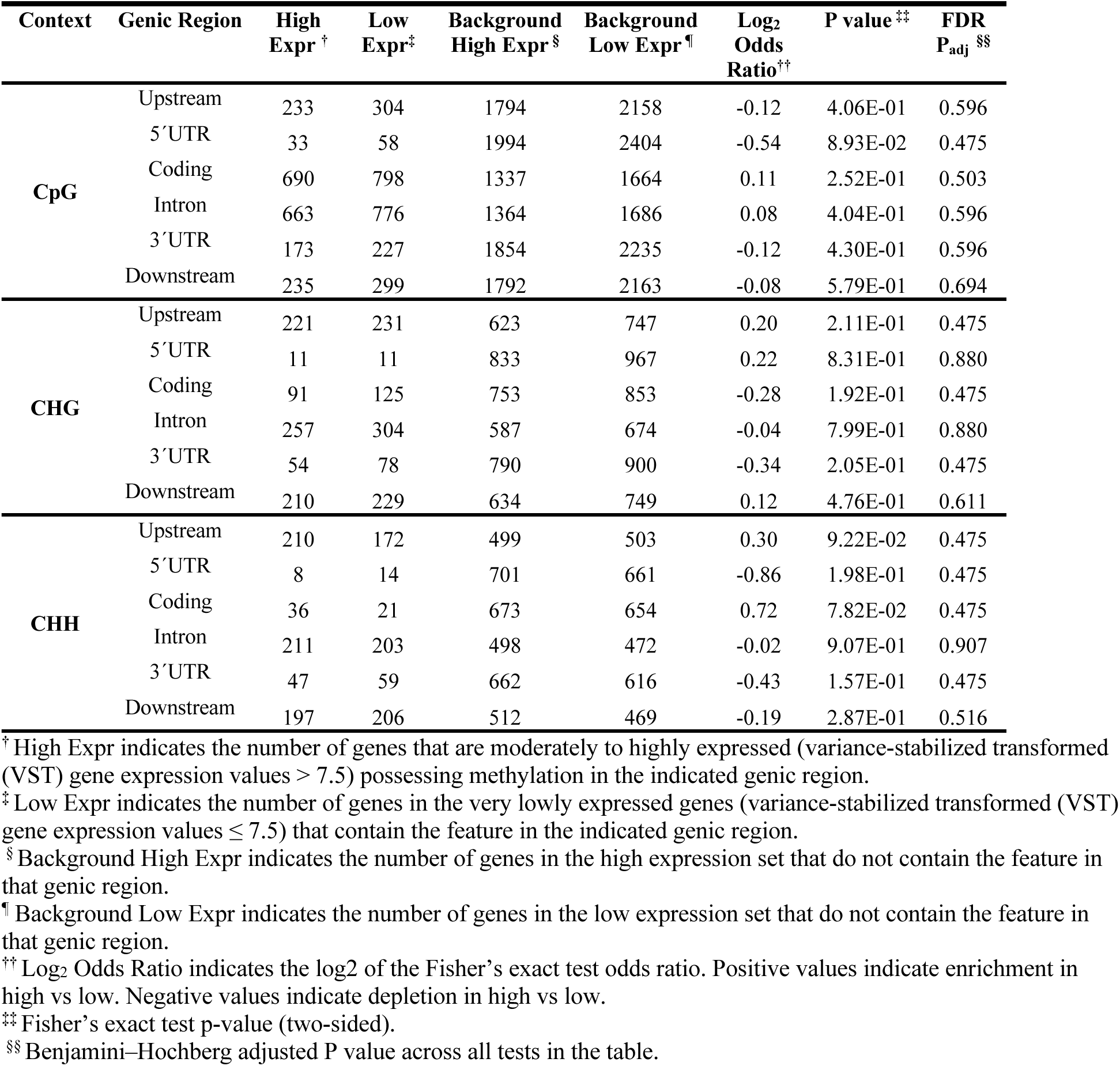
Associations between DNA methylation context and significantly correlated gene expression in different genic regions for Non-TE genes.

**Table S8:**
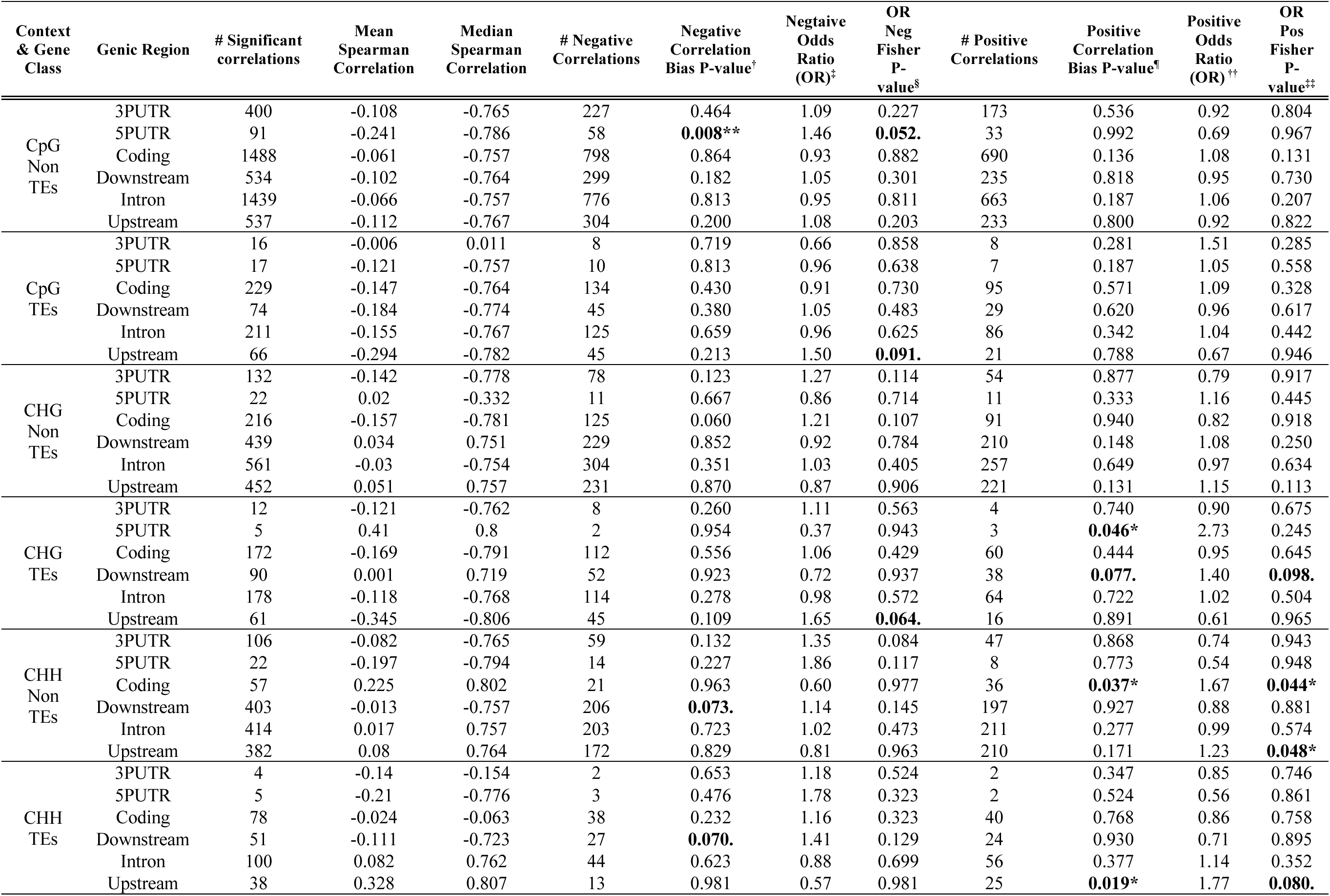
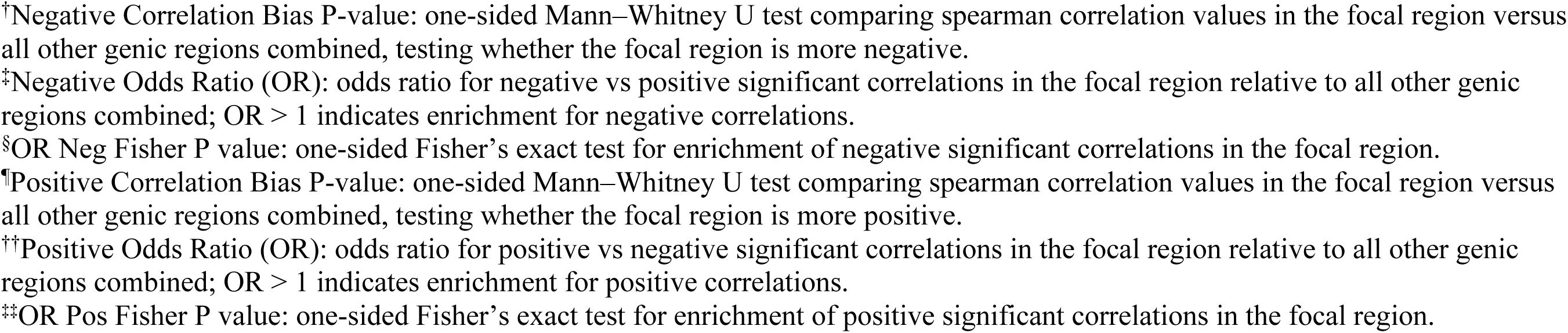
Comparison of Significant Methylation–Expression Correlations by Genic Region in Non-TE and TE associated elements. P-values that are statistically significant are highlighted as bold and annotated as ** < 0.01, * < 0.05,. < 0.1.

| Context & Gene Class | Genic Region | # Significant correlations | Mean Spearman Correlation | Median Spearman Correlation | # Negative Correlations | Negative Correlation Bias P-value<sup>†</sup> | Negative Odds Ratio (OR)<sup>‡</sup> | OR Neg Fisher P-value<sup>§</sup> | # Positive Correlations | Positive Correlation Bias P-value<sup>¶</sup> | Positive Odds Ratio (OR)<sup>††</sup> | OR Pos Fisher P-value<sup>‡‡</sup> |
| --- | --- | --- | --- | --- | --- | --- | --- | --- | --- | --- | --- | --- |
| CpG Non TEs | 3PUTR | 400 | -0.108 | -0.765 | 227 | 0.464 | 1.09 | 0.227 | 173 | 0.536 | 0.92 | 0.804 |
|  | 5PUTR | 91 | -0.241 | -0.786 | 58 | **0.008\*\*** | 1.46 | **0.052.** | 33 | 0.992 | 0.69 | 0.967 |
|  | Coding | 1488 | -0.061 | -0.757 | 798 | 0.864 | 0.93 | 0.882 | 690 | 0.136 | 1.08 | 0.131 |
|  | Downstream | 534 | -0.102 | -0.764 | 299 | 0.182 | 1.05 | 0.301 | 235 | 0.818 | 0.95 | 0.730 |
|  | Intron | 1439 | -0.066 | -0.757 | 776 | 0.813 | 0.95 | 0.811 | 663 | 0.187 | 1.06 | 0.207 |
|  | Upstream | 537 | -0.112 | -0.767 | 304 | 0.200 | 1.08 | 0.203 | 233 | 0.800 | 0.92 | 0.822 |
| CpG TEs | 3PUTR | 16 | -0.006 | 0.011 | 8 | 0.719 | 0.66 | 0.858 | 8 | 0.281 | 1.51 | 0.285 |
|  | 5PUTR | 17 | -0.121 | -0.757 | 10 | 0.813 | 0.96 | 0.638 | 7 | 0.187 | 1.05 | 0.558 |
|  | Coding | 229 | -0.147 | -0.764 | 134 | 0.430 | 0.91 | 0.730 | 95 | 0.571 | 1.09 | 0.328 |
|  | Downstream | 74 | -0.184 | -0.774 | 45 | 0.380 | 1.05 | 0.483 | 29 | 0.620 | 0.96 | 0.617 |
|  | Intron | 211 | -0.155 | -0.767 | 125 | 0.659 | 0.96 | 0.625 | 86 | 0.342 | 1.04 | 0.442 |
|  | Upstream | 66 | -0.294 | -0.782 | 45 | 0.213 | 1.50 | **0.091.** | 21 | 0.788 | 0.67 | 0.946 |
| CHG Non TEs | 3PUTR | 132 | -0.142 | -0.778 | 78 | 0.123 | 1.27 | 0.114 | 54 | 0.877 | 0.79 | 0.917 |
|  | 5PUTR | 22 | 0.02 | -0.332 | 11 | 0.667 | 0.86 | 0.714 | 11 | 0.333 | 1.16 | 0.445 |
|  | Coding | 216 | -0.157 | -0.781 | 125 | 0.060 | 1.21 | 0.107 | 91 | 0.940 | 0.82 | 0.918 |
|  | Downstream | 439 | 0.034 | 0.751 | 229 | 0.852 | 0.92 | 0.784 | 210 | 0.148 | 1.08 | 0.250 |
|  | Intron | 561 | -0.03 | -0.754 | 304 | 0.351 | 1.03 | 0.405 | 257 | 0.649 | 0.97 | 0.634 |
|  | Upstream | 452 | 0.051 | 0.757 | 231 | 0.870 | 0.87 | 0.906 | 221 | 0.131 | 1.15 | 0.113 |
| CHG TEs | 3PUTR | 12 | -0.121 | -0.762 | 8 | 0.260 | 1.11 | 0.563 | 4 | 0.740 | 0.90 | 0.675 |
|  | 5PUTR | 5 | 0.41 | 0.8 | 2 | 0.954 | 0.37 | 0.943 | 3 | **0.046\*** | 2.73 | 0.245 |
|  | Coding | 172 | -0.169 | -0.791 | 112 | 0.556 | 1.06 | 0.429 | 60 | 0.444 | 0.95 | 0.645 |
|  | Downstream | 90 | 0.001 | 0.719 | 52 | 0.923 | 0.72 | 0.937 | 38 | **0.077.** | 1.40 | **0.098.** |
|  | Intron | 178 | -0.118 | -0.768 | 114 | 0.278 | 0.98 | 0.572 | 64 | 0.722 | 1.02 | 0.504 |
|  | Upstream | 61 | -0.345 | -0.806 | 45 | 0.109 | 1.65 | **0.064.** | 16 | 0.891 | 0.61 | 0.965 |
| CHH Non TEs | 3PUTR | 106 | -0.082 | -0.765 | 59 | 0.132 | 1.35 | 0.084 | 47 | 0.868 | 0.74 | 0.943 |
|  | 5PUTR | 22 | -0.197 | -0.794 | 14 | 0.227 | 1.86 | 0.117 | 8 | 0.773 | 0.54 | 0.948 |
|  | Coding | 57 | 0.225 | 0.802 | 21 | 0.963 | 0.60 | 0.977 | 36 | **0.037\*** | 1.67 | **0.044\*** |
|  | Downstream | 403 | -0.013 | -0.757 | 206 | **0.073.** | 1.14 | 0.145 | 197 | 0.927 | 0.88 | 0.881 |
|  | Intron | 414 | 0.017 | 0.757 | 203 | 0.723 | 1.02 | 0.473 | 211 | 0.277 | 0.99 | 0.574 |
|  | Upstream | 382 | 0.08 | 0.764 | 172 | 0.829 | 0.81 | 0.963 | 210 | 0.171 | 1.23 | **0.048\*** |
| CHH TEs | 3PUTR | 4 | -0.14 | -0.154 | 2 | 0.653 | 1.18 | 0.524 | 2 | 0.347 | 0.85 | 0.746 |
|  | 5PUTR | 5 | -0.21 | -0.776 | 3 | 0.476 | 1.78 | 0.323 | 2 | 0.524 | 0.56 | 0.861 |
|  | Coding | 78 | -0.024 | -0.063 | 38 | 0.232 | 1.16 | 0.323 | 40 | 0.768 | 0.86 | 0.758 |
|  | Downstream | 51 | -0.111 | -0.723 | 27 | **0.070.** | 1.41 | 0.129 | 24 | 0.930 | 0.71 | 0.895 |
|  | Intron | 100 | 0.082 | 0.762 | 44 | 0.623 | 0.88 | 0.699 | 56 | 0.377 | 1.14 | 0.352 |
|  | Upstream | 38 | 0.328 | 0.807 | 13 | 0.981 | 0.57 | 0.981 | 25 | **0.019\*** | 1.77 | **0.080.** |

<sup>†</sup>Negative Correlation Bias P-value: one-sided Mann–Whitney U test comparing spearman correlation values in the focal region versus all other genic regions combined, testing whether the focal region is more negative.
<sup>‡</sup>Negative Odds Ratio (OR): odds ratio for negative vs positive significant correlations in the focal region relative to all other genic regions combined; OR > 1 indicates enrichment for negative correlations.
<sup>§</sup>OR Neg Fisher P value: one-sided Fisher’s exact test for enrichment of negative significant correlations in the focal region.
<sup>¶</sup>Positive Correlation Bias P-value: one-sided Mann–Whitney U test comparing spearman correlation values in the focal region versus all other genic regions combined, testing whether the focal region is more positive.
<sup>††</sup>Positive Odds Ratio (OR): odds ratio for positive vs negative significant correlations in the focal region relative to all other genic regions combined; OR > 1 indicates enrichment for positive correlations.
<sup>‡‡</sup>OR Pos Fisher P value: one-sided Fisher’s exact test for enrichment of positive significant correlations in the focal region.

**Table S9:**
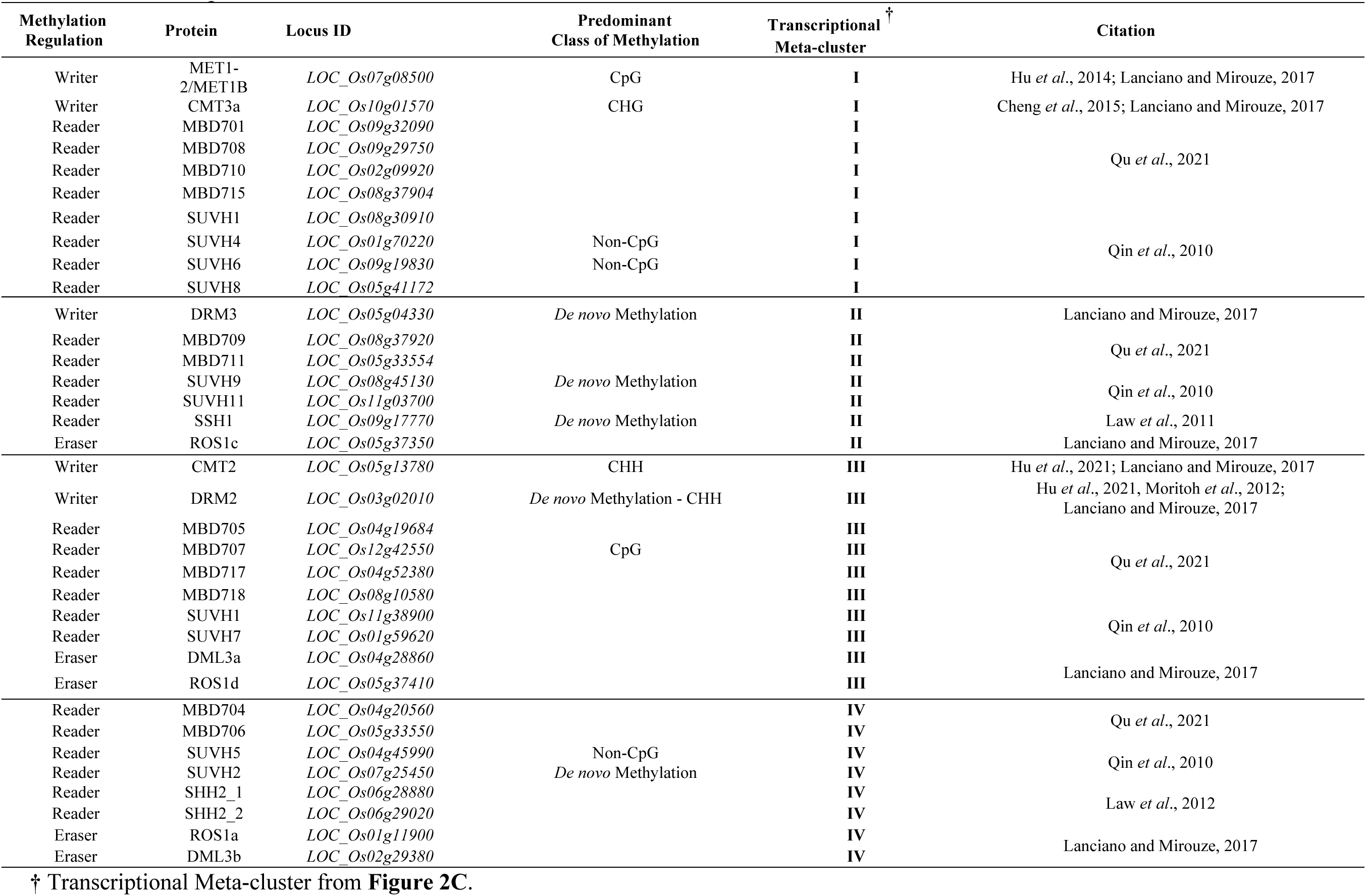
Key Rice Proteins Involved in DNA Methylation Regulation: Classification by Function, Predominant Methylation Class, Internode transcriptional Meta-cluster.

**Figure S8:**
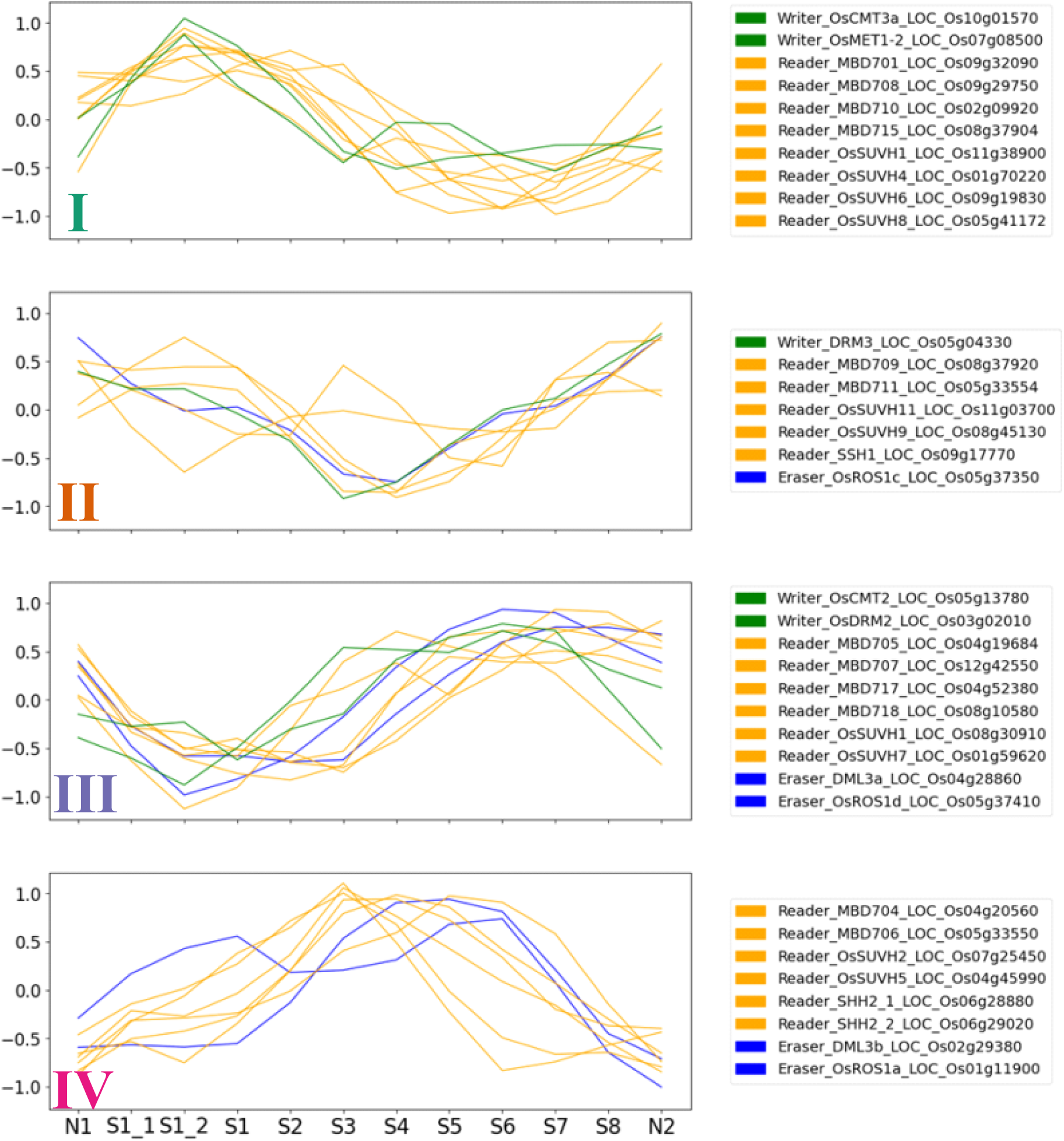
Expression Profiles of DNA Methylation Writers, Readers, and Erasers Across Rice Internode Developmental States. Standardized expression profiles of DNA methylation-related genes categorized as writers (green), readers (orange), and erasers (blue), within each mega cluster (I–IV) across developmental stages (Figure 1D). The y-axis represents standardized expression values (range: -1 to 1) relative to the dataset mean. The x-axis represents the progression of internode developmental states. This analysis highlights the temporal dynamics and clustering of DNA methylation-related genes, revealing distinct functional roles during rice internode development.

**Figure S9:**
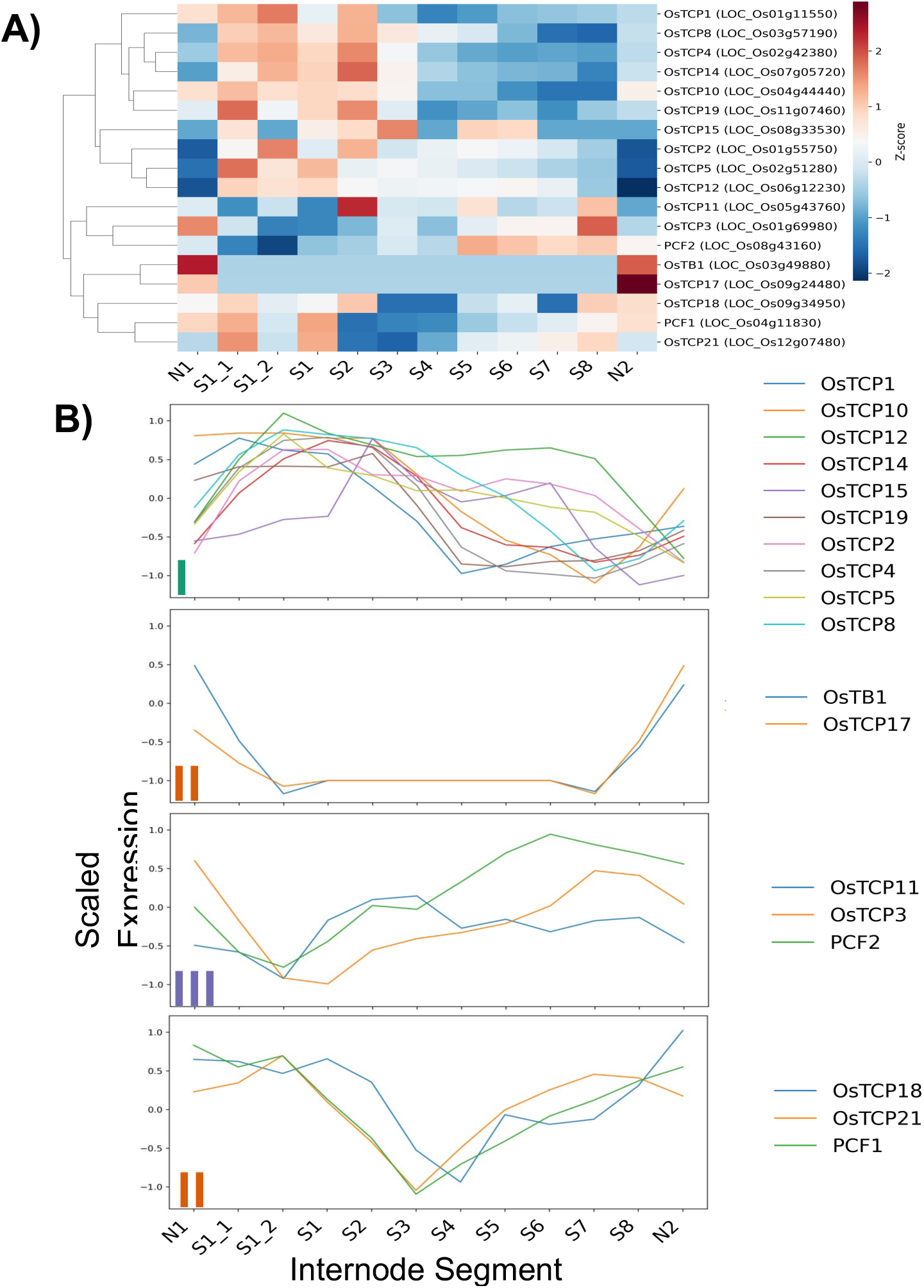
Expression patterns of TCP transcription factors across rice internode development. **(A)** Heatmap showing scaled expression (Z-scores) of TCP transcription factor genes across developmental stages of the rice internode. Rows represent individual TCP genes and columns represent developmental segments. Hierarchical clustering groups genes with similar expression profiles across development. Red indicates higher relative expression, whereas blue indicates lower relative expression. (**B)** Line plots showing expression dynamics of TCP genes grouped into meta-clusters (as defined in Figure 2) based on their developmental profiles. Each line represents the scaled expression of an individual gene across the internode gradient. Meta-clusters I–III are represented, whereas the meta-cluster IV pattern is not observed among TCP genes. These patterns highlight stage-specific regulation of TCP transcription factors during internode growth.

**Figure S10:**
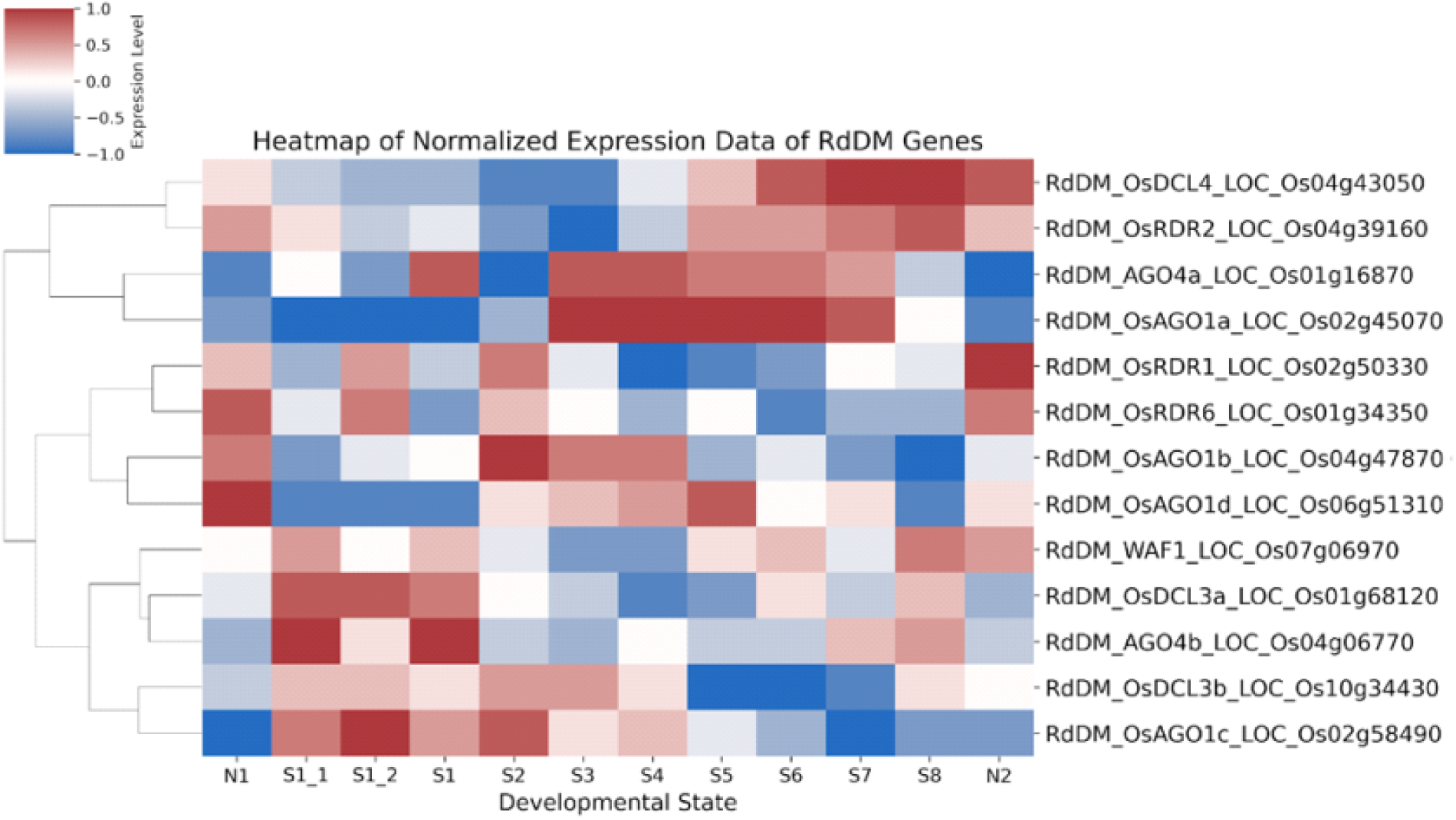
Heatmap of Normalized Expression Data of RdDM Genes Across Developmental States. This heatmap illustrates the normalized expression profiles of RNA-directed DNA methylation (RdDM) pathway genes across rice internode developmental states. Gene expression levels were standardized to a range of -1 (low expression, blue) to 1 (high expression, red). Hierarchical clustering was performed on the rows (genes) to group genes with similar expression patterns. The dendrogram on the left represents the similarity between genes based on their expression profiles. Each row corresponds to a specific RdDM gene, with gene identifiers labeled on the right.

**Figure S11:**
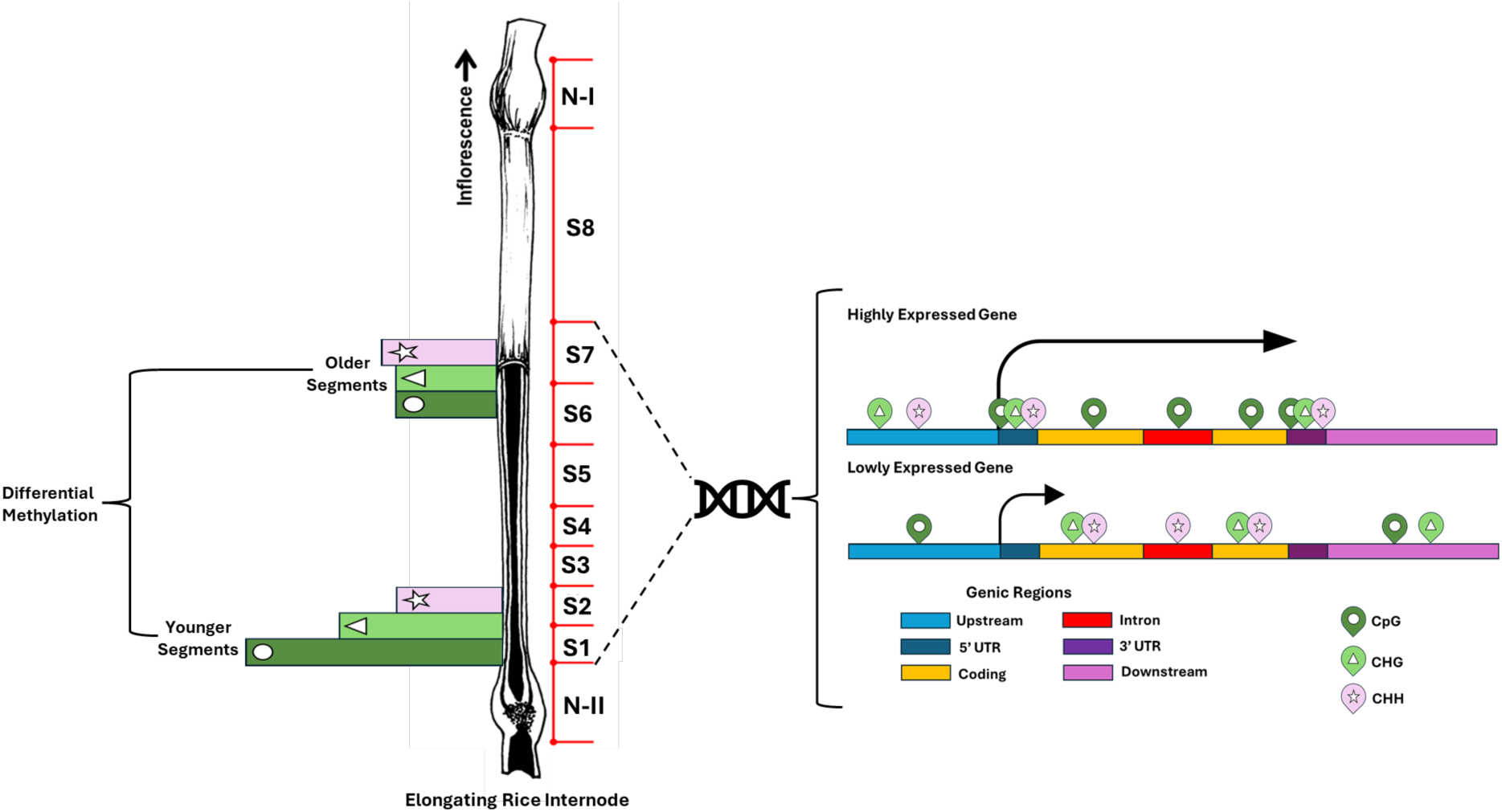
Model illustrating DNA methylation dynamics along the elongating rice internode. A schematic summarizing DNA methylation patterns across internode segments and their relationships with gene expression. Differential methylation in younger segments is predominantly observed in CpG (Green Bar, Circle) and CHG (Light Green, Triangle) contexts, consistent with maintenance methylation, whereas CHH (Pink, Star) methylation occurs across both young and mature segments. A summary schematic highlights methylation contexts and genic regions associated with high and low expression. Higher gene expression was associated with CHH and CHG methylation in upstream regions, CpG methylation within gene bodies, and methylation in 5′ and 3′ untranslated regions. While very low expression was associated with CpG methylation upstream, CHG and CHH methylation within gene bodies, and CpG and CHG methylation downstream.

## Supporting Data

***Data S1:*** Transcriptional meta-clusters: gene membership and functional enrichments

***Data S2:*** Differential methylation between older and younger segments: genes and functional enrichments

***Data S3:*** Genes with significant positive or negative methylation-expression correlations

***Data S4:*** Functional enrichments of genes with significant positive or negative methylation-expression correlations

***Data S5:*** Methylation–transcript correlation results for literature-curated phenylpropanoid, lignin, and flavonoid/tricin-related genes

***Data S6:*** NCBI Sequence Read Archive accession numbers for RNA-seq and bisulfite sequencing datasets

## References

Acemel RD, Maeso I, Gómez-Skarmeta JL. 2017. Topologically associated domains: a successful scaffold for the evolution of gene regulation in animals. WIREs Developmental Biology 6(3): e265.

Akdemir KC, Le VT, Chandran S, Li Y, Verhaak RG, Beroukhim R, Campbell PJ, Chin L, Dixon JR, Futreal PA, et al. 2020. Disruption of chromatin folding domains by somatic genomic rearrangements in human cancer. Nature Genetics 52(3): 294–305.

Bartels A, Han Q, Nair P, Stacey L, Gaynier H, Mosley M, Huang QQ, Pearson JK, Hsieh TF, An YC, et al. 2018. Dynamic DNA Methylation in Plant Growth and Development. Int J Mol Sci 19(7).

Berdasco M, Alcázar R, García-Ortiz MV, Ballestar E, Fernández AF, Roldán-Arjona T, Tiburcio AF, Altabella T, Buisine N, Quesneville H, et al. 2008. Promoter DNA Hypermethylation and Gene Repression in Undifferentiated Arabidopsis Cells. PLoS One 3(10): e3306.

Bewick AJ, Schmitz RJ. 2017. Gene body DNA methylation in plants. Current Opinion in Plant Biology 36: 103–110.

Bray NL, Pimentel H, Melsted P, Pachter L. 2016. Near-optimal probabilistic RNA-seq quantification. Nature Biotechnology 34(5): 525–527.

Candaele J, Demuynck K, Mosoti D, Beemster GTS, Inzé D, Nelissen H. 2014. Differential Methylation during Maize Leaf Growth Targets Developmentally Regulated Genes Plant Physiology 164(3): 1350–1364.

Chen X, Vega-Sanchez ME, Verhertbruggen Y, Chiniquy D, Canlas PE, Fagerstrom A, Prak L, Christensen U, Oikawa A, Chern M, et al. 2013. Inactivation of OsIRX10 leads to decreased xylan content in rice culm cell walls and improved biomass saccharification. Mol Plant 6(2): 570–573.

Cheng C, Tarutani Y, Miyao A, Ito T, Yamazaki M, Sakai H, Fukai E, Hirochika H. 2015. Loss of function mutations in the rice chromomethylase OsCMT3a cause a burst of transposition. The Plant Journal 83(6): 1069–1081.

Domb K, Wang N, Hummel G, Liu C. 2022. Spatial Features and Functional Implications of Plant 3D Genome Organization. Annu Rev Plant Biol 73: 173–200.

Garg R, Narayana Chevala VVS, Shankar R, Jain M. 2015. Divergent DNA methylation patterns associated with gene expression in rice cultivars with contrasting drought and salinity stress response. Scientific Reports 5(1): 14922.

Gaur VS, Channappa G, Chakraborti M, Sharma TR, Mondal TK. 2020. ‘Green revolution’ dwarf gene sd1 of rice has gigantic impact. Briefings in Functional Genomics 19(5-6): 390–409.

Golicz AA, Bhalla PL, Edwards D, Singh MB. 2020. Rice 3D chromatin structure correlates with sequence variation and meiotic recombination rate. Communications Biology 3(1): 235.

Guerriero G, Hausman JF, Ezcurra I. 2015. WD40-Repeat Proteins in Plant Cell Wall Formation: Current Evidence and Research Prospects. Front Plant Sci 6: 1112.

Gui J, Shen J, Li L. 2011. Functional characterization of evolutionarily divergent 4-coumarate:coenzyme a ligases in rice. Plant Physiol 157(2): 574–586.

Hadish JA, Biggs TD, Shealy BT, Bender MR, McKnight CB, Wytko C, Smith MC, Feltus FA, Honaas L, Ficklin SP. 2022. GEMmaker: process massive RNA-seq datasets on heterogeneous computational infrastructure. BMC Bioinformatics 23(1): 156.

Han Z, Crisp PA, Stelpflug S, Kaeppler SM, Li Q, Springer NM. 2018. Heritable Epigenomic Changes to the Maize Methylome Resulting from Tissue Culture. Genetics 209(4): 983–995.

Héberlé É, Bardet AF. 2019. Sensitivity of transcription factors to DNA methylation. Essays Biochem 63(6): 727–741.

Hirano K, Aya K, Morinaka Y, Nagamatsu S, Sato Y, Antonio BA, Namiki N, Nagamura Y, Matsuoka M. 2013. Survey of Genes Involved in Rice Secondary Cell Wall Formation Through a Co-Expression Network. Plant and Cell Physiology 54(11): 1803–1821.

Hu D, Yu Y, Wang C, Long Y, Liu Y, Feng L, Lu D, Liu B, Jia J, Xia R, et al. 2021. Multiplex CRISPR-Cas9 editing of DNA methyltransferases in rice uncovers a class of non-CG methylation specific for GC-rich regions. Plant Cell 33(9): 2950–2964.

Hu L, Li N, Xu C, Zhong S, Lin X, Yang J, Zhou T, Yuliang A, Wu Y, Chen YR, et al. 2014. Mutation of a major CG methylase in rice causes genome-wide hypomethylation, dysregulated genome expression, and seedling lethality. Proc Natl Acad Sci U S A 111(29): 10642–10647.

Ikram AU, Zhang F, Xu Z, Li E, Xue G, Wang S, Zhang C, Yang Y, Su Y, Ding Y. 2022. Chromatin remodeling factors OsYAF9 and OsSWC4 interact to promote internode elongation in rice. Plant Physiology 188(4): 2199–2214.

Irshad F, Li C, Wu HY, Yan Y, Xu JH. 2022. The Function of DNA Demethylase Gene ROS1a Null Mutant on Seed Development in Rice (Oryza sativa) Using the CRISPR/CAS9 System. Int J Mol Sci 23(12).

Jain BP, Pandey S. 2018. WD40 Repeat Proteins: Signalling Scaffold with Diverse Functions. Protein J 37(5): 391–406.

Ji Y, Wang A. 2023. Recent advances in epigenetic triggering of climacteric fruit ripening. Plant Physiology 192(3): 1711–1717.

Jullien PE, Kinoshita T, Ohad N, Berger Fdr. 2006. Maintenance of DNA Methylation during the Arabidopsis Life Cycle Is Essential for Parental Imprinting. The Plant Cell 18(6): 1360–1372.

Kashiwagi T, Togawa E, Hirotsu N, Ishimaru K. 2008. Improvement of lodging resistance with QTLs for stem diameter in rice (Oryza sativa L.). Theoretical and Applied Genetics 117(5): 749–757.

Kawahara Y, de la Bastide M, Hamilton JP, Kanamori H, McCombie WR, Ouyang S, Schwartz DC, Tanaka T, Wu J, Zhou S, et al. 2013. Improvement of the Oryza sativa Nipponbare reference genome using next generation sequence and optical map data. Rice 6(1): 4.

Kenchanmane Raju Sunil K, Ritter Eleanore J, Niederhuth Chad E. 2019. Establishment, maintenance, and biological roles of non-CG methylation in plants. Essays in Biochemistry 63(6): 743–755.

Krueger F, Andrews SR. 2011. Bismark: a flexible aligner and methylation caller for Bisulfite-Seq applications. Bioinformatics 27(11): 1571–1572.

Kumar S, Mohapatra T. 2021. Dynamics of DNA Methylation and Its Functions in Plant Growth and Development. Frontiers in Plant Science 12.

Lam PY, Lui ACW, Yamamura M, Wang L, Takeda Y, Suzuki S, Liu H, Zhu F-Y, Chen M-X, Zhang J, et al. 2019. Recruitment of specific flavonoid B-ring hydroxylases for two independent biosynthesis pathways of flavone-derived metabolites in grasses. New Phytologist 223(1): 204–219.

Lanciano S, Mirouze M. 2017. DNA Methylation in Rice and Relevance for Breeding. Epigenomes 1(2): 10.

Law JA, Vashisht AA, Wohlschlegel JA, Jacobsen SE. 2011. SHH1, a Homeodomain Protein Required for DNA Methylation, As Well As RDR2, RDM4, and Chromatin Remodeling Factors, Associate with RNA Polymerase IV. PLOS Genetics 7(7): e1002195.

Le TM, Tran UPN, Duong YHP, Nguyen KT, Tran VT, Le PK. 2022. Development of a paddy-based biorefinery approach toward improvement of biomass utilization for more bioproducts. Chemosphere 289: 133249.

Lee J, Lee S, Park K, Shin SY, Frost JM, Hsieh PH, Shin C, Fischer RL, Hsieh TF, Choi Y. 2023. Distinct regulatory pathways contribute to dynamic CHH methylation patterns in transposable elements throughout Arabidopsis embryogenesis. Front Plant Sci 14: 1204279.

Li W, Chen G, Xiao G, Zhu S, Zhou N, Zhu P, Zhang Q, Hu T. 2020. Overexpression of TCP transcription factor OsPCF7 improves agronomic trait in rice. Molecular Breeding 40(5): 48.

Li X, Yang Y, Yao J, Chen G, Li X, Zhang Q, Wu C. 2009. FLEXIBLE CULM 1 encoding a cinnamyl-alcohol dehydrogenase controls culm mechanical strength in rice. Plant Mol Biol 69(6): 685–697.

Liang W, Li J, Sun L, Liu Y, Lan Z, Qian W. 2022. Deciphering the synergistic and redundant roles of CG and non-CG DNA methylation in plant development and transposable element silencing. New Phytol 233(2): 722–737.

Lin F, Williams BJ, Thangella PAV, Ladak A, Schepmoes AA, Olivos HJ, Zhao K, Callister SJ, Bartley LE. 2017. Proteomics Coupled with Metabolite and Cell Wall Profiling Reveal Metabolic Processes of a Developing Rice Stem Internode. Frontiers in Plant Science 8.

Lister R, O’Malley RC, Tonti-Filippini J, Gregory BD, Berry CC, Millar AH, Ecker JR. 2008. Highly Integrated Single-Base Resolution Maps of the Epigenome in *Arabidopsis*. Cell 133(3): 523–536.

Liu C, Cheng Y-J, Wang J-W, Weigel D. 2017. Prominent topologically associated domains differentiate global chromatin packing in rice from Arabidopsis. Nature Plants 3(9): 742–748.

Lou S, Lee H-M, Qin H, Li J-W, Gao Z, Liu X, Chan LL, Kl Lam V, So W-Y, Wang Y, et al. 2014. Whole-genome bisulfite sequencing of multiple individuals reveals complementary roles of promoter and gene body methylation in transcriptional regulation. Genome Biology 15(7): 408.

Margueron R, Justin N, Ohno K, Sharpe ML, Son J, Drury Iii WJ, Voigt P, Martin SR, Taylor WR, De Marco V, et al. 2009. Role of the polycomb protein EED in the propagation of repressive histone marks. Nature 461(7265): 762–767.

Markulin L, Škiljaica A, Tokić M, Jagić M, Vuk T, Bauer N, Leljak Levanić D. 2021. Taking the Wheel – de novo DNA Methylation as a Driving Force of Plant Embryonic Development. Frontiers in Plant Science 12.

Martin GT, Seymour DK, Gaut BS. 2021. CHH Methylation Islands: A Nonconserved Feature of Grass Genomes That Is Positively Associated with Transposable Elements but Negatively Associated with Gene-Body Methylation. Genome Biology and Evolution 13(8): evab144.

McArthur E, Capra JA. 2021. Topologically associating domain boundaries that are stable across diverse cell types are evolutionarily constrained and enriched for heritability. Am J Hum Genet 108(2): 269–283.

Moritoh S, Eun C-H, Ono A, Asao H, Okano Y, Yamaguchi K, Shimatani Z, Koizumi A, Terada R. 2012. Targeted disruption of an orthologue of DOMAINS REARRANGED METHYLASE 2, OsDRM2, impairs the growth of rice plants by abnormal DNA methylation. The Plant Journal 71(1): 85–98.

Muyle AM, Seymour DK, Lv Y, Huettel B, Gaut BS. 2022. Gene Body Methylation in Plants: Mechanisms, Functions, and Important Implications for Understanding Evolutionary Processes. Genome Biology and Evolution 14(4): evac038.

Nazipova A, Gorshkov O, Eneyskaya E, Petrova N, Kulminskaya A, Gorshkova T, Kozlova L. 2022. Forgotten Actors: Glycoside Hydrolases During Elongation Growth of Maize Primary Root. Frontiers in Plant Science 12.

Ni P, Huang N, Nie F, Zhang J, Zhang Z, Wu B, Bai L, Liu W, Xiao C-L, Luo F, et al. 2021. Genome-wide detection of cytosine methylations in plant from Nanopore data using deep learning. Nature Communications 12(1): 5976.

Patil V, McDermott HI, McAllister T, Cummins M, Silva JC, Mollison E, Meikle R, Morris J, Hedley PE, Waugh R, et al. 2019. APETALA2 control of barley internode elongation. Development 146(11).

Pikaard CS, Mittelsten Scheid O. 2014. Epigenetic regulation in plants. Cold Spring Harb Perspect Biol 6(12): a019315.

Qin F-J, Sun Q-W, Huang L-M, Chen X-S, Zhou D-X. 2010. Rice SUVH Histone Methyltransferase Genes Display Specific Functions in Chromatin Modification and Retrotransposon Repression. Molecular Plant 3(4): 773–782.

Qu M, Zhang Z, Liang T, Niu P, Wu M, Chi W, Chen ZQ, Chen ZJ, Zhang S, Chen S. 2021. Overexpression of a methyl-CpG-binding protein gene OsMBD707 leads to larger tiller angles and reduced photoperiod sensitivity in rice. BMC Plant Biol 21(1): 100.

Ruan C, Yang J, Yang S, Huang W, Zhou Y, Wang P, Yu D. 2026. Class I TCP transcription factor OsTCP4 suppresses plant height via negatively regulating the green revolution gene SD1 (OsGA20ox2) in rice (Oryza sativa). Plant Diversity.

Santos AP, Gaudin V, Mozgová I, Pontvianne F, Schubert D, Tek AL, Dvořáčková M, Liu C, Fransz P, Rosa S, et al. 2020. Tidying-up the plant nuclear space: domains, functions, and dynamics. Journal of Experimental Botany 71(17): 5160–5178.

Secco D, Wang C, Shou H, Schultz MD, Chiarenza S, Nussaume L, Ecker JR, Whelan J, Lister R. 2015. Stress induced gene expression drives transient DNA methylation changes at adjacent repetitive elements. eLife 4: e09343.

Stefansson OA, Sigurpalsdottir BD, Rognvaldsson S, Halldorsson GH, Juliusson K, Sveinbjornsson G, Gunnarsson B, Beyter D, Jonsson H, Gudjonsson SA, et al. 2024. The correlation between CpG methylation and gene expression is driven by sequence variants. Nature Genetics 56(8): 1624–1631.

Stirnimann CU, Petsalaki E, Russell RB, Müller CW. 2010. WD40 proteins propel cellular networks. Trends Biochem Sci 35(10): 565–574.

Sullivan AM, Arsovski AA, Lempe J, Bubb KL, Weirauch MT, Sabo PJ, Sandstrom R, Thurman RE, Neph S, Reynolds AP, et al. 2014. Mapping and dynamics of regulatory DNA and transcription factor networks in A. thaliana. Cell Rep 8(6): 2015–2030.

Sun Y, Dong L, Zhang Y, Lin D, Xu W, Ke C, Han L, Deng L, Li G, Jackson D, et al. 2020. 3D genome architecture coordinates trans and cis regulation of differentially expressed ear and tassel genes in maize. Genome Biology 21(1): 143.

Takeda Y, Koshiba T, Tobimatsu Y, Suzuki S, Murakami S, Yamamura M, Rahman MM, Takano T, Hattori T, Sakamoto M, et al. 2017. Regulation of CONIFERALDEHYDE 5-HYDROXYLASE expression to modulate cell wall lignin structure in rice. Planta 246(2): 337–349.

Tan F, Zhou C, Zhou Q, Zhou S, Yang W, Zhao Y, Li G, Zhou DX. 2016. Analysis of Chromatin Regulators Reveals Specific Features of Rice DNA Methylation Pathways. Plant Physiol 171(3): 2041–2054.

Tirot L, Jullien PE, Ingouff M. 2021. Evolution of CG Methylation Maintenance Machinery in Plants. Epigenomes DOI: 10.3390/epigenomes5030019

Tsuda K, Maeno A, Nonomura KI. 2023. Heat shock-inducible clonal analysis reveals the stepwise establishment of cell fate in the rice stem. Plant Cell 35(12): 4366–4382.

Valton AL, Dekker J. 2016. TAD disruption as oncogenic driver. Curr Opin Genet Dev 36: 34–40.

Wang B, Smith SM, Li J. 2018. Genetic Regulation of Shoot Architecture. Annual Review of Plant Biology 69(Volume 69, 2018): 437–468.

Wang G, Li H, Meng S, Yang J, Ye N, Zhang J. 2020. Analysis of Global Methylome and Gene Expression during Carbon Reserve Mobilization in Stems under Soil Drying Plant Physiol 183(4): 1809–1824.

Wang J, Qi M, Liu J, Zhang Y. 2015. CARMO: a comprehensive annotation platform for functional exploration of rice multi-omics data. Plant J 83(2): 359–374.

Wang W, Qin Q, Sun F, Wang Y, Xu D, Li Z, Fu B. 2016. Genome-Wide Differences in DNA Methylation Changes in Two Contrasting Rice Genotypes in Response to Drought Conditions. Frontiers in Plant Science 7.

Wang Z, Baulcombe DC. 2020. Transposon age and non-CG methylation. Nature Communications 11(1): 1221.

Wei L, Gu L, Song X, Cui X, Lu Z, Zhou M, Wang L, Hu F, Zhai J, Meyers BC, et al. 2014. Dicer-like 3 produces transposable element-associated 24-nt siRNAs that control agricultural traits in rice. Proceedings of the National Academy of Sciences 111(10): 3877–3882.

Wreczycka K, Gosdschan A, Yusuf D, Grüning B, Assenov Y, Akalin A. 2017. Strategies for analyzing bisulfite sequencing data. Journal of Biotechnology 261: 105–115.

Xie L, Wen D, Wu C, Zhang C. 2022. Transcriptome analysis reveals the mechanism of internode development affecting maize stalk strength. BMC Plant Biology 22(1): 49.

Yamauchi T, Johzuka-Hisatomi Y, Terada R, Nakamura I, Iida S. 2014. The MET1b gene encoding a maintenance DNA methyltransferase is indispensable for normal development in rice. Plant Molecular Biology 85(3): 219–232.

Yang XC, Hwa CM. 2008. Genetic modification of plant architecture and variety improvement in rice. Heredity 101(5): 396–404.

Yano K, Ookawa T, Aya K, Ochiai Y, Hirasawa T, Ebitani T, Takarada T, Yano M, Yamamoto T, Fukuoka S, et al. 2015. Isolation of a novel lodging resistance QTL gene involved in strigolactone signaling and its pyramiding with a QTL gene involved in another mechanism. Mol Plant 8(2): 303–314.

Yao X, Ma H, Wang J, Zhang D. 2007. Genome-Wide Comparative Analysis and Expression Pattern of TCP Gene Families in Arabidopsis thaliana and Oryza sativa. Journal of Integrative Plant Biology 49(6): 885–897.

Yin M, Wang S, Wang Y, Wei R, Liang Y, Zuo L, Huo M, Huang Z, Lang J, Zhao X, et al. 2024. Impact of Abiotic Stress on Rice and the Role of DNA Methylation in Stress Response Mechanisms. Plants 13(19): 2700.

Zhang H, Lang Z, Zhu J-K. 2018. Dynamics and function of DNA methylation in plants. Nature Reviews Molecular Cell Biology 19(8): 489–506.

Zhang H, Zheng R, Wang Y, Zhang Y, Hong P, Fang Y, Li G, Fang Y. 2019. The effects of Arabidopsis genome duplication on the chromatin organization and transcriptional regulation. Nucleic Acids Res 47(15): 7857–7869.

Zhang K, Mosch K, Fischle W, Grewal SIS. 2008. Roles of the Clr4 methyltransferase complex in nucleation, spreading and maintenance of heterochromatin. Nature Structural & Molecular Biology 15(4): 381–388.

Zhang X, Yazaki J, Sundaresan A, Cokus S, Chan SW, Chen H, Henderson IR, Shinn P, Pellegrini M, Jacobsen SE, et al. 2006. Genome-wide high-resolution mapping and functional analysis of DNA methylation in arabidopsis. Cell 126(6): 1189–1201.

Zheng K, Wang L, Zeng L, Xu D, Guo Z, Gao X, Yang D-L. 2021. The effect of RNA polymerase V on 24-nt siRNA accumulation depends on DNA methylation contexts and histone modifications in rice. Proceedings of the National Academy of Sciences 118(30): e2100709118.

Zhong Z, Feng S, Duttke SH, Potok ME, Zhang Y, Gallego-Bartolomé J, Liu W, Jacobsen SE. 2021. DNA methylation-linked chromatin accessibility affects genomic architecture in *Arabidopsis*. Proceedings of the National Academy of Sciences 118(5): e2023347118.

